# Intrinsic disorder in elicitin-like effectors: Molecular shields in the arms race of biotrophic pathogens

**DOI:** 10.64898/2026.04.09.717489

**Authors:** Monja Schmid, Daniel Gómez-Pérez, Samuel Quinzer, Edda von Roepenack-Lahaye, Ariane Kemen, Eric Kemen

**Author notes:** Author for correspondence: Ariane Kemen, Eric Kemen.

## Abstract

Plant pathogens establish colonization through effector secretion to modulate host immune responses. Within bacterial effectors intrinsically disordered regions (IDRs) contribute to diverse functions and pathogenicity, but their role in effectors of phytopathogenic oomycetes remains poorly understood. Here, analysis of oomycete secretomes across lifestyles reveal widespread intrinsic disorder in secreted proteins, particularly in apoplastic elicitin and elicitin-like effectors (ELLs). Combining *in vitro* and *in planta* experiments, we show that IDRs in ELLs from the biotrophic pathogen *Albugo candida* reduce immune recognition, consistent with an IDR-mediated shielding mechanism. This function is transferable, as fusion of an ELL-derived IDR to the immunogenic elicitin INF1 from *Phytophthora infestans* abolishes its recognition. These findings identify intrinsic disorder as a functional feature of apoplastic effectors that modulates host detection and promotes pathogen fitness.

**Graphical summary:** 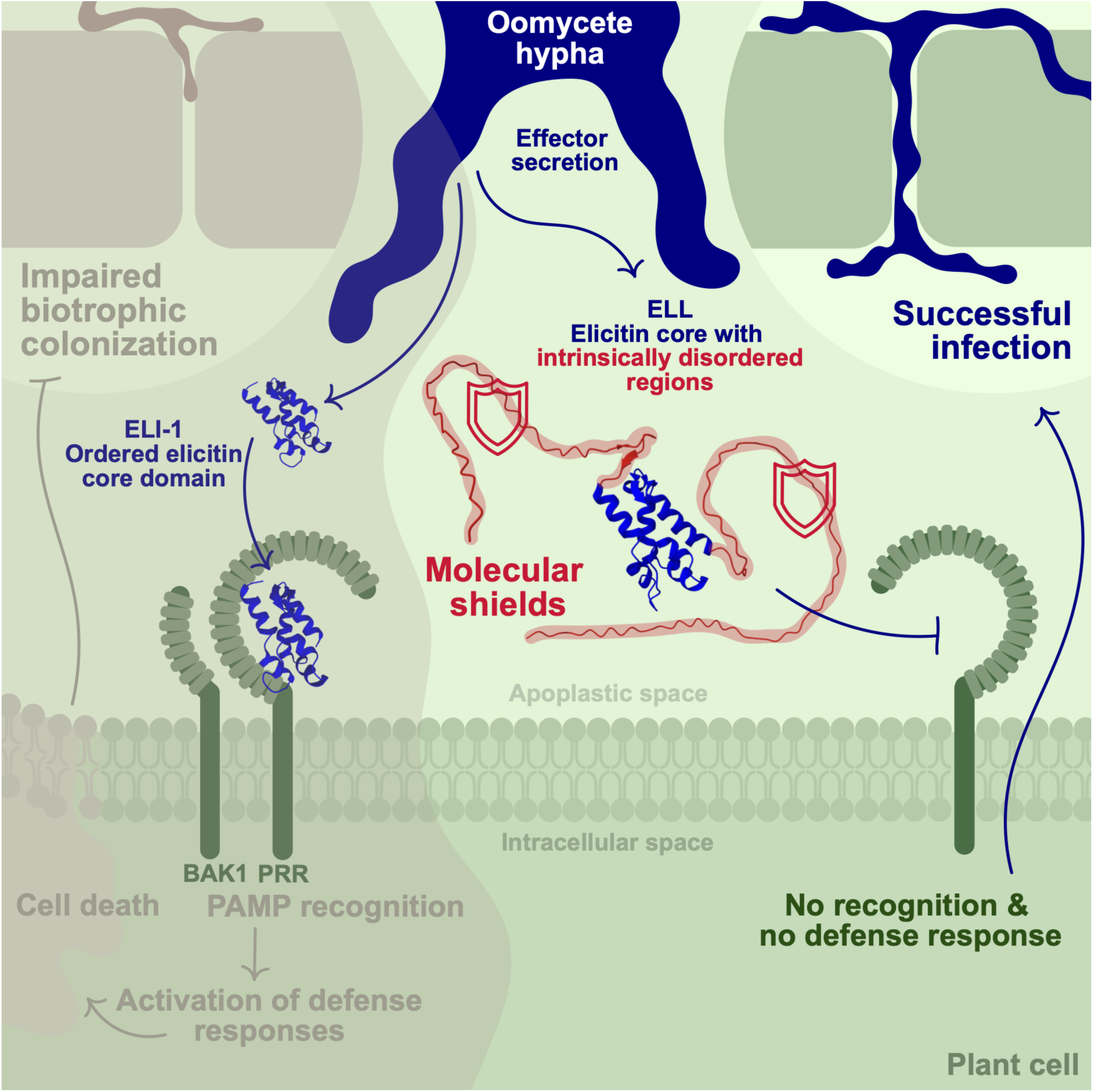

## Introduction

At the core of plant–pathogen interactions lies a complex evolutionary cycle in which microbial effectors and plant resistance constantly co-evolve. Pattern-triggered immunity (PTI) is one such defense mechanism, consisting of pattern recognition receptors (PRRs) on the plant cell surface that detect pathogen-associated molecular patterns (PAMPs), secreted by the pathogen to promote infection. Strategies for evading recognition or countering the induced defenses determine pathogen colonization success and shape the outcomes in this molecular arms race^1–4^.

For many proteins involved in pathogenesis and defense, structural features can be reliably derived from the amino acid sequence^5,6^. However, deriving function from structure is often not feasible. In this context, intrinsically disordered regions (IDRs) within proteins add to the challenge by defying the classical key-lock protein model^7,8^, leading to the consideration of proteins as multifunctional entities and replacing the one-structure-one-function model with a structure-function-continuum^9–12^. Protein IDRs lack fixed folding leading to an unstable three-dimensional structure. This state allows for increased post-translational modifications, exposure of linear motives for binding partners, cleavage by proteases and accelerated evolution^13^. IDRs exhibit a compositional bias toward charged, polar amino acids and often lack bulky or hydrophobic residues^14^. Structural flexibility in secreted proteins facilitates their translocation and offers a strategy to control their activity by folding in a context or stimulus-dependent manner^15^. In addition to intracellular roles^16^, proteins with IDRs govern interactions between plant hosts and microbial colonizers, including pathogens. For example, IDRs facilitate protein aggregation into robust amyloid fibrils that frequently act as scaffolds in host-associated biofilms^17–19^. They confer several advantages to plant colonizers, contributing to processes ranging from adhesion and surface hydrophobicity to signaling and symbiosis^20,21^. IDRs can also contribute to microbial interactions through inhibitory activities, for example by enabling targeted modification of plant-associated microbial communities^22^. Additionally, IDRs may conceal pathogenic bacterial proteins from recognition by the plant, potentially acting as molecular shields that reduce binding of effector to immune receptors such as R proteins^23^. Remarkably, although abundant in nature, IDRs have hardly been studied experimentally in this context, leaving their contribution to interactions between effectors and plant immunity largely unexplored.

Despite discoveries regarding effector evolution and the functionality of IDRs in fungal and bacterial pathogens^5,15,24^, research on phytopathogenic oomycetes remains limited. Of particular interest in the context of host-pathogen interactions are hemibiotrophic and obligate biotrophic oomycetes due to their intimate adaptation to host plants^4^. Hemibiotrophs such as *Phytophthora* species typically repress defenses and modulate immune responses during an early biotrophic phase of infection, before switching to a necrotrophic mode in which they kill the host and feed on decaying tissues^25^. In contrast, obligate biotrophs strictly rely on living plant tissue to proliferate, with adaptations that have led to reduced selection for maintenance of certain biosynthetic pathways and loss of secondary metabolism^26–28^. Members of this lifestyle include *Albugo candida* and *Albugo laibachii*^29–31^, both infecting the model plant *Arabidopsis thaliana* with *A. candida* exhibiting an extended host range^32^. Upon infection, both pathogens cause white blister rust disease^33^. Secreted effectors and membrane-anchored proteins play key roles in the infection process by ensuring nutrient uptake, colonization and microbial dominance^22,34,35^. While some immunity strategies of *A. thaliana* against *Albugo* pathogens have been described^36,37^, the exact mechanisms behind many protein-based interactions remain poorly understood.

Apoplastic elicitin proteins, unique to oomycetes, have been studied for several decades in the context of the molecular arms race^38,39^. Their ability to bind extracellular sterols makes them plausible effectors, given that most oomycetes are unable to synthesize sterols *de novo*^40^. However, beyond correlations between elicitin expression and sterol availability, many details of sterol recruitment remain elusive^41–43^. Better characterized is their role as PAMPs, inducing basal defense, hypersensitive response, necrosis and cell death upon recognition by dedicated PRRs^44,45^. These include the two leucine-rich repeat receptor-like proteins (LRR-RLPs) ELR (elicitin response) from *Solanum microdontum*^46^ and REL (responsive to elicitin) from *Nicotiana benthamiana*^47^. Both recognize INF1, a well-studied elicitin from *Phytophthora infestans*^48^, belonging to class 1. Four classes of elicitins have been described, all sharing a conserved core domain and secretion signal. In addition to conventional secretion, some can be covalently bound to the cell wall^49^. Class 1 elicitins (ELI-1) contain only the conserved core, whereas classes 2, 3 and 4 (ELI-2/3/4) possess additional domains of varying length and amino acid composition. Compared to elicitins, elicitin-like proteins (ELLs) show variations in both the core and additional domains. The conserved core domain is 98 amino acids in length and contains a characteristic cysteine pattern^50^. Research has primarily focused on ELI-1 from *Phytophthora* species, leaving the functions of additional domains in other classes and ELLs largely unexplored. Some studies suggest roles in membrane attachment via glycosylphosphatidylinositol (GPI) anchoring or glycosylation, but their extended functionality remains poorly understood^51,52^.

In this study, we investigate whether the functions attributed to IDRs in bacterial effectors are likewise relevant in ELL effectors from the oomycete pathogen *A. candida*. We first assess the prevalence of IDRs in secreted proteins of phytopathogenic oomycetes, revealing a widespread enrichment of disorder in effector repertoires. We demonstrate that IDRs in ELLs can function as molecular shields that contribute to evasion of host immunity. Consistent with this, IDRs in *Albugo* ELLs facilitate reduced recognition by BAK1-dependent PRRs in *A. thaliana* and promote amyloid formation, with the shielding function also being effective in other hosts. Our findings highlight intrinsic disorder as a key determinant of pathogenic strategies and host interactions in oomycetes, particularly in pathogens with an obligate biotrophic lifestyle.

## Methods and Materials

### Computational analysis of oomycete secretomes

We compiled the proteomes of available oomycete species^53^ and used SignalP6.0^54^ with the parameters Eukarya and fast to predict the classical secretome (<u>Supp. Table 1</u>). For all proteins identified as having a secretion signal, we performed intrinsic disorder predictions using flDPnn^55^. The predictor assigns a score to each amino acid in the sequence. An amino acid is considered to be disordered when the score is equal or higher than 0.3, as suggested by the developers. A significant intrinsically disordered domain within a protein has more than 15 consecutive disordered amino acids in agreement with other publications^56,57^. We calculated a total disorder score for each protein by summing the length of all significant disordered domains and dividing it by the total length of the protein. We performed the disorder prediction on the full-length proteins including the secretion signal, but we did not include the area of the secretion signal into the calculation of the total disorder score. Proteins with a disorder score of 0 were considered to be completely ordered, proteins with a score higher than 0 were considered to contain intrinsically disordered regions (IDRs). We also annotated the lifestyle of each of the oomycete species based on literature consensus^53^: obligate biotrophic, hemibiotrophic, necrotrophic or other. The category other included all lifestyles that were not plant pathogenic and organisms for which the lifestyle has not yet been clearly defined. We analyzed functional descriptions of proteins with disorder scores of 0.2 or higher (disordered secretome) and with scores lower than 0.2 (ordered secretome). For this, we used InterProScan 5^58^ and compiled recurring functional terms.

### Computational analysis of proteins with annotated elicitin domains

We used the InterPro database to compile all proteins with an annotated elicitin domain (IPR002200)^59^. We considered all proteins that were annotated prior to 30.01.2024. We created a table including the respective organism for each protein and the host and lifestyle category for each organism (<u>Supp. Table 2</u>). Lifestyles included obligate biotrophic, hemibiotrophic, necrotrophic or other as described in the previous section. Host categories included plant, animal, fungus/oomycete, none for saprotrophs and unknown for all organisms with unknown hosts. Lifestyle and host classification were based on a previous publication^53^ and literature consensus. Proteins for which no exact species could be assigned were not included as the lifestyle cannot be reliably determined for these organisms. We used the dataset of oomycete proteomes mentioned above to correlate the count of elicitins per species with the respective overall protein count. For prediction of secretion signal sequences, we used SignalP6.0^54^ with the parameters Eukarya and fast. We performed an intrinsic disorder prediction on the amino acid sequences of each protein using flDPnn^55^.

To group the elicitins by class, we followed the guidelines from Janků et al.^50^ for classification. We considered the classical elicitin core domain as (1) being annotated as elicitin domain in the InterPro database, (2) having a length between 96 and 100 amino acids and (3) having six cysteine residues. Proteins consisting of just the core domain with no additional domains were considered traditional class 1 elicitins (ELI-1), proteins with the core domain and additional domains class 2/3/4 elicitins (ELI-2/3/4). Proteins with an annotated elicitin domain but differences in core and additional domains were labeled elicitin-like proteins (ELLs).

For predicting amyloid forming properties, we used AMYPred-FRL^60^ on the sequences without secretion signal. Each protein received a score by the predictor. We considered a protein as amyloid forming when the score was equal or higher than 0.5, as recommended by the developers. All prediction scores can be found in <u>Supp. Table 2</u>.

For creating the sequence similarity tree of *Albugo candida* and *Albugo laibachii* elicitins, we used the R packages ape, msa and phangorn. For prediction of glycosylphosphatidylinositol (GPI) anchoring we used NetGPI 1.1^61^. For structure predictions we used AlphaFold3^62^ and for alignments, the RCSB structural alignment tool^63^.

### Plant growth and infection with *Albugo candida* and *Albugo laibachii*

We grew *Arabidopsis thaliana* plants of ecotype WS-0 and infected them with *A. candida* strain Nc2 or *A. laibachii* strain Nc14 as described before^22^. Briefly, the plants were inoculated by spraying a water-spore suspension onto the leaves and keeping them at high humidity for 48 hours before transferring them to short day growth conditions (8 h light at 21°C, 16 h dark at 16°C).

### Expression of elicitin-like candidates from *Albugo candida* and *Albugo laibachii*

In order to confirm expression of the proteins annotated with an elicitin domain in *A. candida* and *A. laibachii*, we isolated RNA from *A. thaliana* plants infected with either of the strains at different timepoints after infection. For *A. candida*, we isolated RNA during the earlier phase of infection prior to the formation of spores at five days post infection (dpi) and during the late stage (14 dpi), characterized by heavy spore formation and breaching of the epidermis. For *A. laibachii* we isolated RNA during the middle stage just after the appearance of the first pustules (8 dpi) and additionally during the late stage (14 dpi). For the extraction, we used the RNeasy Plant Mini Kit (Qiagen, Cat. No. 74904) followed by treatment with the TURBO DNA free Kit (Invitrogen, ThermoFisher, Cat. No. 2238G). For the synthesis of complementary DNA (cDNA) we used the SuperScript IV Reverse Transcriptase (ThermoFisher, Cat. No. 18090010) with Murine RNAse inhibitor (NewEngland Biolabs, Cat No. M0314S) treatment. For *A. candida,* we found 17 proteins annotated with an elicitin domain according to InterPro and for *A. laibachii* nine (<u>Supp. Table 3</u>). We designed primers for each to amplify them from the cDNA (<u>Supp. Table 4</u>). For amplification we used Phusion^®^ High-Fidelity DNA polymerase (NewEngland Biolabs, Cat. No. M0530S) with recommended conditions, followed by visualization with an agarose gel electrophoresis. We extracted the bands from the gel with the Nucleospin Gel and PCR clean-up Kit (Macherey-Nagel, Cat. No. 740609.250) and used the LGC sequencing service for Sanger sequencing to check sequence accuracy.

We additionally checked for the presence of these proteins in the apoplast of *A. thaliana* infected with either *A. candida* Nc2 or *A. laibachii* Nc14 at 10 dpi. For this we used a previously published proteomics dataset^22,64^.

### Cloning and expression of elicitin-like candidates in C-DAG system

For testing the amyloid forming abilities of the ELL candidates, we used the curli-dependent amyloid generator system (C-DAG) with *Escherichia coli* VS45 as heterologous expression host. This strain harbors plasmid pVS76 for overexpression of csgG, relevant for the amyloid export system and inducible with IPTG, and plasmid pExport for the overexpression of the target candidate amyloids and inducible with L-arabinose^65^. For the amplification of the ELL candidates from *A. candida* and *A. laibachii* we used Phusion^®^ High-Fidelity DNA polymerase and cDNA as template. For candidates we could not amplify from the cDNA, we extracted DNA from leaves in the late stage of infection using the Quick-DNA^TM^ Fungal/Bacterial Miniprep Kit (Zymo Research, Cat. No. D6005) and amplified the candidates from the genomic DNA, since none of them had introns. We designed the primers as recommended by the developers of the C-DAG system excluding the secretion signal (<u>Supp. Table 4</u>).

We cloned the ELL candidates into plasmid pExport as suggested by the supplier. In short, we digested the plasmid and the amplified inserts with NotI-HF (NewEngland Biolabs, Cat. No R3189S) and XbaI (NewEngland Biolabs, Cat. No R0145S) and purified them after agarose gel electrophoresis with the Nucleospin Gel and PCR clean-up Kit. We performed ligation with T4 DNA Ligase (NewEngland Biolabs, Cat. No M0202S) and chemically transformed the final plasmids into the VS45. We created partial versions of some of the candidates: E205Core, E206Core, E212Core and E215Core (only containing the predicted elicitin core domains) and E212IDR (only containing the C-terminal IDR of E212).

To assess the amyloid-forming properties of the cloned ELLs, we performed colony-phenotype-assays. We inoculated 10 mL of LB medium (supplemented with 100 µg/mL carbenicillin, 25 µg/mL chloramphenicol and 1 mM IPTG) from overnight cultures to an OD600 of 0.05 followed by incubation for 30 min at 37°C and 200 rpm. We then spotted 8 µL of each culture onto inducing LB agar plates (supplemented with 100 µg/mL carbenicillin, 25 µg/mL chloramphenicol and 1 mM IPTG, 0.2% (w/v) L-arabinose and 10 µg/mL Congo Red) and incubated these for 4 days at 22°C before taking pictures. Due to weak cell growth on the plate after induction of E212IDR, we conducted an assay with a lower concentration of inducer (0.05 % L-arabinose). We included pVS72, the provided positive control plasmid expressing Sup35, a prion from *Saccharomyces cerevisiae* and pVS105, the negative control plasmid expressing Sup35 without the aggregating domain. Background color in the pictures was normalized with Fiji (ImageJ)^66^ and the macros White balance correction_1.0.ijm.

We prepared samples for transmission electron microscopy by lightly touching the colonies from the induction plates with a mesh copper grid followed by brief drying. We performed negative staining with aqueous uracyl acetate incubation for 30-45 seconds and imaged the samples with a JEM-1400 Flash (JEOL, Japan).

### Cloning and expression of elicitin-like candidates in the PURExpress^®^ system

For functional analysis of the ELL candidates, we expressed them via the PURExpress^®^ system (NewEngland Biolabs, Cat. No E6800L). For cloning the target sequences into the provided PURExpress^®^ plasmid we used the In-Fusion^®^ cloning method with the In-Fusion^®^ Snap Assembly Master Mix (Takara Bio, Cat. No 638948). We designed the primers as recommended by the supplier (<u>Supp. Table 5</u>, <u>Supp. Table 6</u>) and amplified the *A. candida* ELL candidates as described in the previous section. For the candidate INF1 from *Phytophthora infestans* we isolated DNA from hyphal structures grown at 16°C on V8 agar plates (30 mM CaCO_3_, 20% [v/v] tomato juice, 1.5% [w/v] agar, filtered through 3 layers of microcloth [25 µm pore size]). We resuspended the hyphae on the plate with cold sterile Milli-Q water, transferred the sample to a tube and cooled it on ice for 45 min. We extracted the DNA with the DNeasy^®^ Plant Mini Kit (Cat. No 69104). Amplification was done with custom primers (<u>Supp. Table 5</u>) and Phusion^®^ High-Fidelity DNA polymerase as for the other candidates.

In order to replace the DHFR control gene with the gene of interest, we digested the PURExpress^®^ DHFR control plasmid with the enzymes NdeI (NewEngland Biolabs, Cat. No R0111S) and XhoI (NewEngland Biolabs, Cat. No R0146S) followed by treatment with the T4 DNA polymerase (NewEngland Biolabs, Cat. No M0203S) to create blunt ends. We performed cloning as suggested by the supplier and transformed *E. coli* OneShot^TM^ Top10 competent cells with the final plasmids. We isolated the plasmids with the NucleoSpin Plasmid isolation Kit (Macherey-Nagel, Cat. No 740588.250) and used these for the expression. Alongside the full ELL candidates E206, E212 and E207, we created partial versions: E206Core and E212Core (containing only the predicted elicitin core domains) and E212IDR (only containing the IDR of E212). We created a hybrid ELL candidate INF1-E212IDR by cloning inf1 into the plasmid already containing E212IDR.

We followed the instructions of the supplier for expression using 500 ng as plasmid template for each 25 µL expression. We also added PURExpress^®^ Disulfide bond Enhancer (NewEngland Biolabs, Cat. No E6820) and 20 units of Murine RNase Inhibitor (NewEngland Biolabs, Cat. No M0314) to the reaction. Incubation time and temperature were 3 h and 37° respectively. We verified successful expression by SDS PAGE and proteomics. For constructs INF1 and INF1-E212IDR, we created versions with a N- or C-terminal HiBit tag and used the NanoGlo^®^ HiBit blotting system (Promega, Cat. No N2410) to detect and visualize the proteins via Western Blot.

Targeted LC–MS/MS analysis was performed on the peptides extracted from the aggregation band of E212 that we observed after SDS PAGE. An in-gel digest was performed according to Shevchenko et al.^67^ with slight modifications (reduction and alkylation of cysteine bridges: 10 mM DTT in 20 mM ammonium bicarbonate, 55 mM iodoacetamide in 20 mM ammonium bicarbonate; protein digest: trypsin (0,02µg/µl, proteomics grade, porcine; Merck) in 40 mM ABC Puffer/9% ACN), 37C ON, final digest volume 53 µl; peptide extraction: 50%ACN/0.1%FA, evaporation in SpeedVac, avoiding complete desiccation; original protease digest volume was finally diluted ½ in acidic water (0.1% formic acid). All solvents used in the peptide profiling analyses were LCMS grade.

The targeted LC-MSMS profiling analysis was performed using a Micro-LC M5 (Trap and Elute) and a QTRAP6500+ (Sciex) operated in MRM mode. Chromatographic separation was achieved on a HaloFused C18 column (150 x 0.5 mm (particle size 2.7 µm; Sciex) and a Luna C18(2) trap column (5 μm; 100 Å; 20×0.5 mm; Phenomenex) with a column temperature of 55 °C. The following binary gradient was applied for the main column at a flow rate of 16 µl min-1: 0 - 6 min, linear from 95% to 70% A; 6 - 15 min, linear from 70% to 50% A; 15 - 18 min, linear 50% to 5% A; 18 – 18.5 min, linear from 5% to 95% A; 18.5 - 20 min, isocratic 95% A (A: water, 0.1% aq. formic acid; B: acetonitrile, 0.1% aq. formic acid). The samples were concentrated on the trap column using the following conditions: flow rate 25 µl min-1: 0 - 2.7 min isocratic 95% A; at 2.5 min start main gradient. The injection volume was 50 μl. Analytes were ionized using an Optiflow Turbo V ion source equipped with a SteadySpray T micro electrode in positive ion mode (ion spray voltage: 4800 V). Following additional instrument settings were applied: nebuliser and heater gas, nitrogen, 25 and 45 psi; curtain gas, nitrogen, 30 psi; collision gas, nitrogen, medium; source temperature, 200 °C; entrance potential, +10 V; collision cell exit potential, +10V. The dwell time for all MRMs was 5 msec. The declustering potential was kept at 80 V.

All MRM information was calculated using the Skyline 23.1^68^ (specific peptide settings: missed cleavage1; Carbamidomethyl (C)). Spectra were acquired and sequence coverage analyzed using the vendors software Analyst v1.7.3 and ProteinPilot™ v5.0.2 with the Paragon™ algorithm (v5.0.2.0.5174), using a custom FASTA database consisting of the *A. candida* proteome (Proteome ID: UP000053237, UniProt download: 16.11.2023), the proteolytic enzyme and the target protein E212. All monitored transitions are listed in <u>Supp. Table 7</u>. Homology searches were conducted using BLASTP 2.17.0+ against the NCBI NR protein database (accessed on 18.11.2023)^69^.

### Infiltration analysis of expressed ELL candidates in *Arabidopsis thaliana*

For infiltration assays, we grew *A. thaliana* plants for around eight weeks until they reached sufficient size for infiltration. We injured the surface of the bottom side of the leaves, once on the right side and once on the left, with a needle and used a 1 mL syringe to press the protein solution into the leaf. The infiltration was repeated on the following day. The proteins were expressed directly prior to each infiltration and diluted after expression 1:300 in 10 mM MgCl_2_. The proteins were expressed directly prior to each infiltration. Apart from the expressed ELL candidates we included a negative control containing all the PURExpress^®^ components but no expressed protein. When not taken out for infiltration, the plants were kept at short-day growth conditions (8 h light at 21°C, 16 h dark at 16°C). On day six after the first infiltration, we screened for necrosis and cell death in the infiltrated leaves. To do so, we punched out three leaf disks (5 mm diameter) on each side from the infiltrated area around the puncture location and floated all six leaf disks together in 600 µL sterile milliQ water in a 100-well plate. After one hour, we discarded the water to remove electrolytes leaking from the cut edges and replaced it with 1200 µL fresh sterile milliQ water. We used a conductometer with electrodes for each of the 100-wells and measured conductivity every 20 minutes for ten hours. We excluded the first 90 minutes from consideration to ensure sufficient leakage of all samples prior to comparison.

We created three sets including different treatments: set 1 included infiltrations with E212, E212Core, E212IDR and buffer; set 2 included E206, E206Core, E207 and a buffer control. For these two sets we used *A. thaliana* ecotype WS-0. In set 3, we only used E212Core and buffer to infiltrate ecotype Col-0, bak1-3 (SALK_034523) and bak1-5^70^. The sets were handled in independent experimental setups. For each set, we performed three separate rounds, identical in the setup but with independently grown plants. We used 4-5 plants per candidate in each round and infiltrated 4-5 leaves per plant, leading to 20 replicates. The buffer control in set 3 was only added in round 3. The experimental overview is summarized in <u>Supp. Table 8</u>. We considered each round as a biological replicate and performed statistics on the conductivity measurements accordingly. For this, we first used the R package lme4 to perform the linear mixed-effects model analysis with the protein treatments as fixed effects and the rounds or biological replicates as random effects. We then used package emmeans to perform a post-hoc test for pairwise comparisons with Tukey adjustments.

### Infiltration analysis of expressed elicitin-like candidates in *Nicotiana benthamiana*

We conducted the infiltration of *N. benthamiana* leaves in the same fashion as described for *A. thaliana*, with one exception. Depending on the size of the leaves, we infiltrated several spots on the leaf, rather than two as it was done for *A. thaliana*. The plants were grown in the greenhouse under long-day conditions (12 h light, 24 °C, 12 h dark, 22 °C) with relative humidity up to 70% for 4-5 weeks. We distributed the infiltration treatments into four sets. For set 1, we infiltrated INF1, E212Core, E206Core and buffer. For set 2, we infiltrated INF1, INF1-E212IDR and buffer. For this set, we also included completely untreated leaves in the cell death quantification. The proteins were produced directly before infiltration on day 1 and day 2. For set 1 and 2, we infiltrated the same protein on both days. For set 3 and 4, we infiltrated different proteins on day 1 and 2. In set 3, we infiltrated (1) diluted INF1 (half of the regular amount) on both days, (2) E212IDR on day 1 and INF1 on day 2, (3) buffer on day 1 and INF1 on day 2, (4) buffer on both days. For set 4, we infiltrated (1) INF1 on both days, (2) a mix separately containing E212IDR and INF1 on both days, (3) hybrid INF1-E212IDR on both days, (4) E212IDR on both days. The sets were handled in independent experimental setups. For each set we performed three separate rounds, identical in the setup but with independently grown plants. We infiltrated 2-3 leaves of 2-4 plants per treatment and round. The experimental overview is shown in <u>Supp. Table 9</u> and <u>Supp. Figure 1</u>.

For quantifying cell death, we scanned the infiltrated leaves for fluorescence on the fluor stage of a Typhoon FLA 9500 (GE Healthcare) with voltage of the photo-multiplier tube of 450 V and a pixel size of 50-100 µm. The images were processed with Fiji^66^. We used the Circle_Tool.ijm to measure pixel intensity with a radius set to 50-100 pixels around the infiltration point to compare the intensity between the treatments. Statistics were performed with lme4 and emmeans packages as described in the previous section.

## Results

### Intrinsic disorder is a dominant feature of oomycete secretomes

In bacterial effectors, IDRs contribute to translocation, signaling, host mimicry, and evasion of detection, thereby influencing pathogenesis and the molecular arms race^23^. To address the limited understanding of IDR function in secreted effectors of oomycete pathogens, we analyzed the predicted proteomes and secretomes of oomycete species spanning lifestyles from obligate biotrophy to necrotrophy. As reported previously^53^, obligate biotrophic pathogens such as *Albugo* and *Hyaloperonospora* sp. had the smallest proteomes and the smallest classically predicted secretomes relative to their proteome size (<u>Supp. Figure 2</u>; <u>Supp. Table 1</u>). IDR predictions revealed that, across most species and independent of lifestyle, approximately 50% of secreted proteins contain IDRs (Figure 1A). To identify protein groups enriched in IDRs, we annotated secreted proteins with at least 20% of disorder using InterProScan and analyzed recurring functional terms. Across all lifestyles, elicitin effectors appeared most often among IDR-containing proteins, whereas in hemibiotrophic pathogens, RxLR effectors were most prevalent within this group (Figure 1B, <u>Supp. Figure 3</u>). In contrast, secretome fractions lacking intrinsic disorder were enriched in functional enzymes such as kinases, peptidases and glycosyl hydrolases, independent of the lifestyle (<u>Supp. Figure 4</u>). Focusing on obligate biotrophic pathogens, we identified elicitin effectors as well as cysteine-rich secretory protein, antigen 5, and pathogenesis-related 1 (CAP) domain containing proteins as major IDR-associated groups (Figure 1C). CAP domain-containing proteins contribute to plant pathogen virulence^71^. PsCAP1 from the hemibiotroph *Phytophthora sojae* triggers immune responses via recognition by an LRR-RLP, associating with co-receptors BAK1 and SORBIR1^72^. Though, PsCAP1 exhibits little disorder (<u>Supp. Figure 5</u>), in contrast to highly disordered CAP proteins from *A. candida* (<u>Supp. Figure 6</u>).

**Figure 1:**
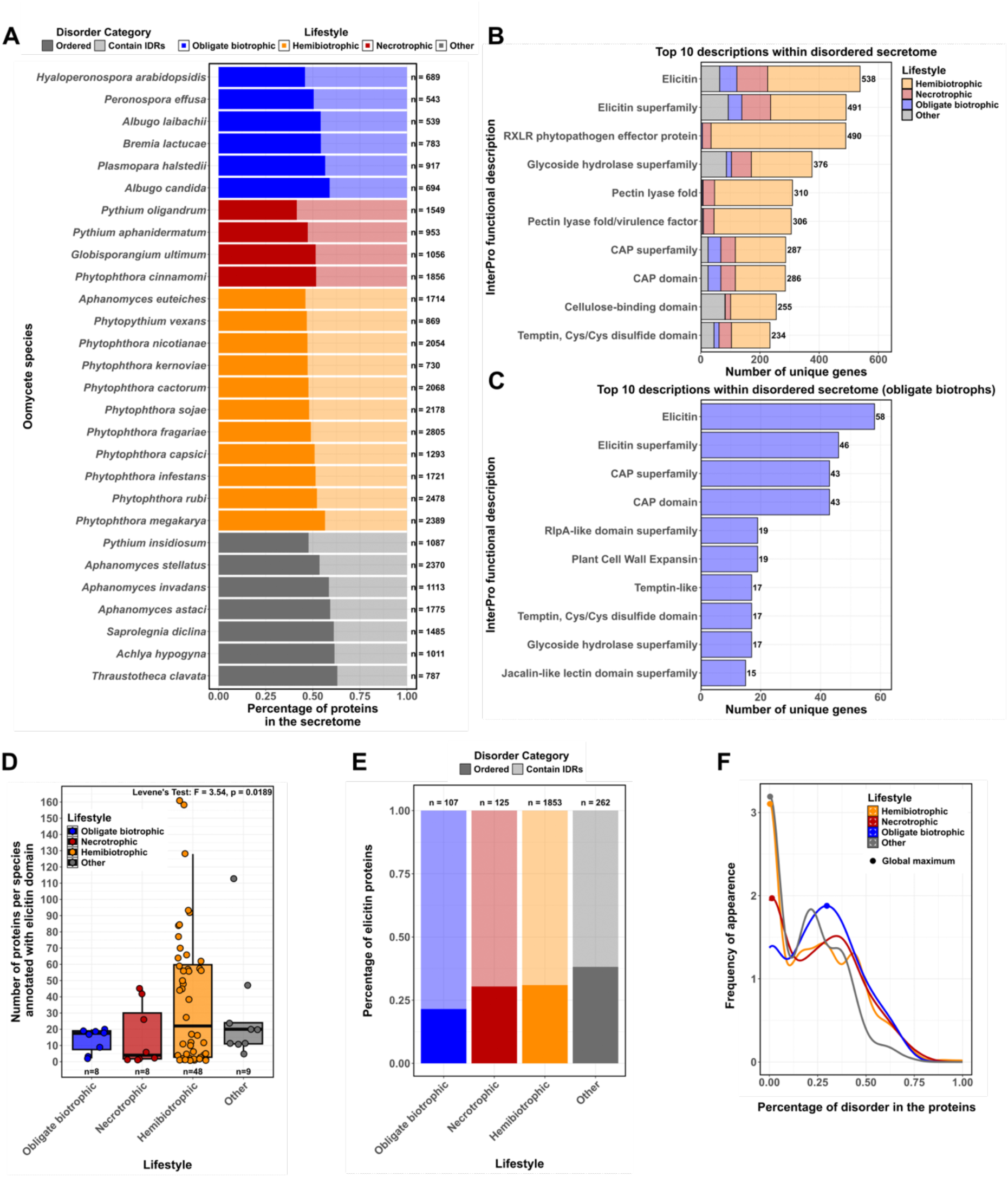
Analysis of intrinsic disorder within the secretomes of oomycete pathogens and within elicitin effectors. **A** Analysis of ordered proteins and proteins containing intrinsically disordered regions (IDRs) within the predicted secretomes of oomycete species. Stacked bar plot showing relative percentage of ordered (dark shade) and IDR containing (light shade) proteins in the secretome per species with the colors representing the different oomycete lifestyles; n indicates the total number of proteins in the respective secretome. SignalP6.0 was used to identify the secretome for each species (<u>Supp. Figure 2</u>, <u>Supp. Table 1</u>), flDPnn was used to predict the disorder scores. **B, C** Occurrence of functional descriptions (InterProScan 5) within the disordered portion of the secretome (disorder score 0.2 and higher) across all oomycete species (B) and across obligate biotrophic species (C). Shown are the ten most frequent terms. The colors represent the lifestyles. Terms occurring within ordered portions and hemibiotrophic species can be found in <u>Supp. Figure 3</u> and <u>Supp. Figure 4</u>. **D** Count of proteins annotated with an elicitin domain per individual oomycete species (strain) plotted by lifestyle (collected from InterPro IPR002200); n describes the number of strains assigned to the specific lifestyle. Levene’s Test was used to assess the differences in the variances of the distributions; the difference is significant when p < 0.05. Counts per individual strain can be found in <u>Supp. Figure 7</u> and <u>Supp. Table 1</u>. The height of the boxes represents the interquartile range (IQR) with the median shown as black horizontal line; the whiskers indicate the range of data within 1.5 times IQR from the quartiles. Correlation to proteome size is shown in <u>Supp. Figure 8</u>. **E** Stacked bar plot showing the relative percentage of ordered (dark) and IDR containing (light) proteins annotated as elicitin per lifestyle; n indicates the total elicitin count of the respective lifestyle. Plot with the exact percentages can be found in <u>Supp. Figure 9</u>. **F** Density curve of percentage of disorder (total disorder score) within elicitin proteins per lifestyle (indicated by the colors). The dot shows the highest peak of each distribution.

Despite their established role in plant-pathogen interactions, little is known about how structural properties of elicitins and ELLs vary across oomycete genera, prompting us to analyze all known elicitin/ELL proteins from oomycetes focusing on the relationship between intrinsic disorder and lifestyle. We compiled a dataset of 2347 proteins annotated with elicitin domains (IPR002200), spanning 63 different oomycete species with 73 individual strains (<u>Supp. Table 2</u>). Although the dataset was biased towards *Phytophthora* species (39 of 63 species) due to genome/proteome data availability, it included 14 additional oomycete genera, ensuring representation of diverse lifestyles. The number of elicitins per strain varied significantly by lifestyle. Hemibiotrophic *Phytophthora* species exhibited the highest counts, led by *Phytophthora fragariae* containing 161 elicitins, whereas obligate biotrophs, led by *Bremia lactucae*, had no more than 20 elicitins (Figure 1D, <u>Supp. Figure 7</u>). Variance analysis (Levene’s test, p < 0.05) confirmed significant differences in elicitin distributions across lifestyles. As already noted, overall predicted proteome size is often linked to lifestyle with biotrophs being at the smallest end^53^. Hence, we studied the link between overall protein and elicitin count per strain and observed a linear correlation with a moderate fit (R^2^ = 0.338, p < 0.001; <u>Supp. Figure 8</u>). This suggests that elicitin abundance is influenced by both lifestyle and overall proteome size.

Additionally, we analyzed the secondary and tertiary structure of elicitins across lifestyles. Notably, 78.5% of elicitins from obligate biotrophs contained IDRs, compared to ∼69% of elicitins from hemibiotrophs and necrotrophs and ∼61.8% from other lifestyles (Figure 1E, <u>Supp. Figure 9</u>). Analyzing density curves of the total disorder score per lifestyle (Figure 1F) revealed a peak at 29.8% protein disorder for obligate biotrophs, whereas necrotrophs, hemibiotrophs, and other lifestyles peaked at substantially lower values (1.4%, 0.4% and 0.3%, respectively). These findings show that IDRs are abundant in elicitin effectors and suggest that intrinsic disorder may represent a hallmark of obligate biotrophy, contributing to infection strategies. To determine the location of IDRs within proteins annotated as elicitins, we first classified them based on their sequence properties. Per species, the majority of them were classified as ELL. Interestingly, obligate biotrophic proteins were exclusively classified as ELI-2/3/4 or ELLs (Figure 2A, <u>Supp. Figure 10</u>, <u>Supp. Table 2</u>). While the elicitin core domain was largely ordered, consistent with a stable three-dimensional structure, IDRs were prevalent in additional domains across species and lifestyle. INF2a from *P. infestans,* for example, contains a disordered C-terminal domain like many other proteins annotated as elicitins in this pathogen (<u>Supp. Figure 11</u>, <u>Supp. Figure 12</u>). Similar examples were found in other hemibiotrophs such as *P. sojae* or the biotroph *Hyaloperonospora arabidopsidis* (<u>Supp. Figure 13</u>, <u>Supp. Figure 14</u>).

**Figure 2:**
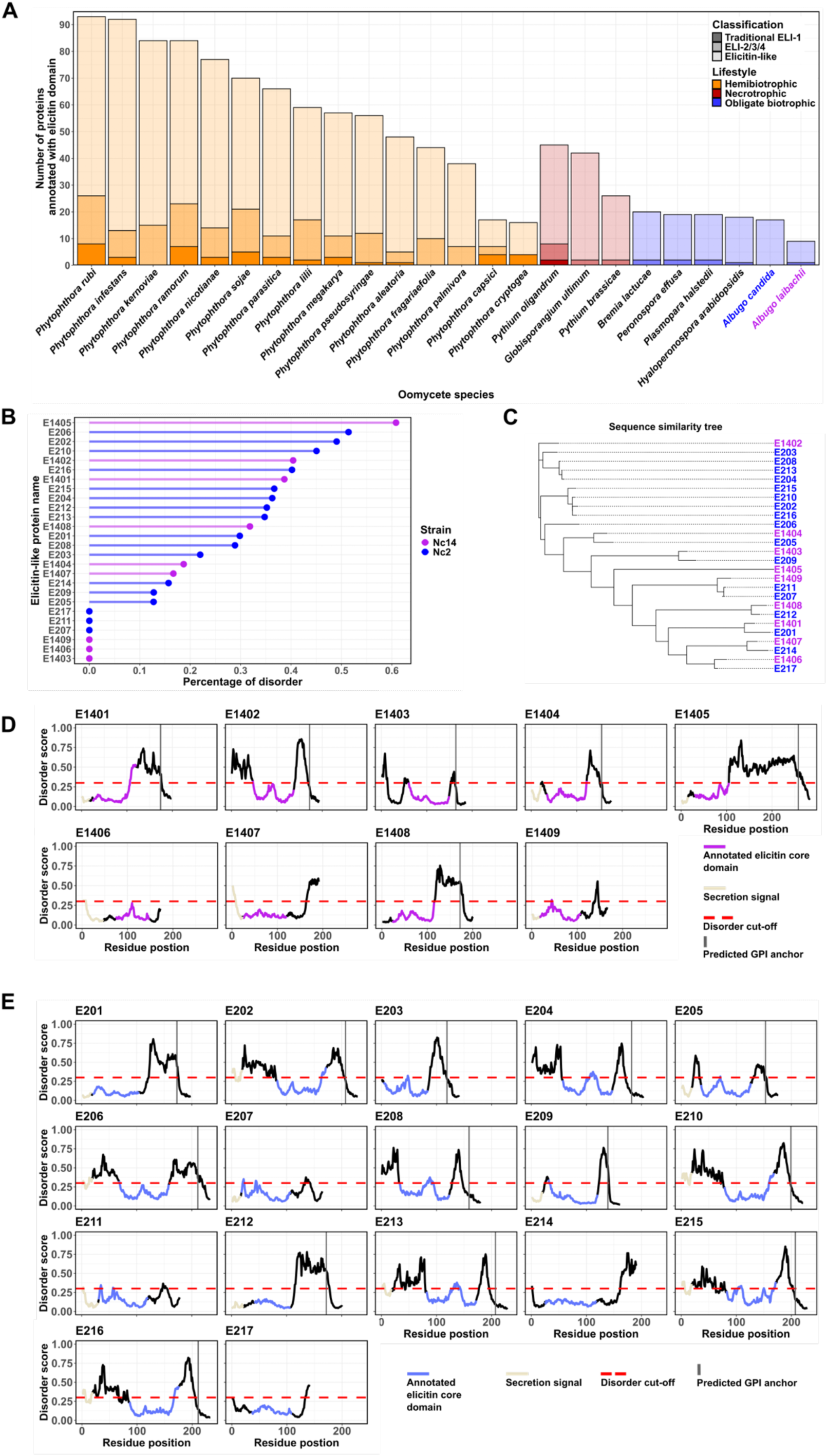
Elicitin characteristics across pathogenic oomycetes and within the obligate biotrophic species *Albugo candida* Nc2 and *Albugo laibachii* Nc14. **A** Classification of proteins with annotated elicitin domain per species. Traditional class 1 elicitins (ELI-1) are defined as having a conserved core domain (96-100 amino acids with six cysteine residues) and no additional domains (dark shade). Class 2/3/4 elicitins (ELI-2/3/4) have the same core domain but exhibit additional domains (mid-tone shade). Elicitin-like proteins (ELLs) have more variable core and appending domains (light shade)^50^. Colors represent the different oomycete lifestyles. The depicted species were chosen as representatives due to their relevance and well annotated proteomes. The remaining species from the dataset can be found in <u>Supp. Figure 10</u>. **B** Percentage of disorder (Total disorder score) per elicitin/ELLs from *Albugo* species. Nc14 and Nc2 ELLs are labeled in magenta and blue, respectively. **C** Phylogenetic tree showing sequence similarities among the *Albugo* ELLs. Magenta shows Nc14, blue shows Nc2. **D, E** Disorder profiles of elicitins from Nc14 (D) and Nc2 (E) depicting disorder scores per amino acid residue in the sequence. Magenta (Nc14) and blue (Nc2) sequence stretch mark annotated elicitin domain, beige stretch mark the secretions signal, the grey vertical line marks the residue of the predicted glycosylphosphatidylinositol (GPI) anchoring site (ω-sites) according to NetGPI 1.1. The red dashed line shows the cut-off for positive disorder predictions according to the flDPnn disorder predictor.

### Intrinsic disorder is prevalent in elicitin-like effectors of *Albugo* pathogens

Building on these findings, we focused on ELLs from two obligate biotrophic pathogens, *A. candida* and *A. laibachii*, comprising 17 and nine ELLs, respectively. Except for six ELLs, expression was verified by amplifying the corresponding genes from cDNA derived from RNA of infected *A. thaliana* leaves. Sequencing revealed slight differences in five ELLs compared to their accession sequences. For E202 and E208, these differences were consistent across independent amplifications, resulting in eleven and three amino acid changes, respectively. Furthermore, we detected four ELLs (E206, E212, E1405, and E1408) in the apoplastic space using a proteomics dataset generated from apoplastic fluid of infected *A. thaliana* plants at ten days post infection (<u>Supp. Table 3</u>)^22,64^. Disorder prediction revealed high prevalence of IDRs in *Albugo* ELLs, consistent with the distribution of disorder observed in obligate biotrophic elicitins and ELLs (Figure 1F, Figure 2B). Interestingly, most ELLs from *A. laibachii* had homologs in *A. candida* based on sequence similarity. *A. candida*, however, contained a diversified cluster of four closely related ELLs similar in sequence to E1402, as well as a cluster with no homology to any *A. laibachii* ELL (Figure 2C). Regarding IDR localization, *A. laibachii* ELLs typically contain a C-terminal IDR, whereas about half of the *A. candida* ELLs contain both C-terminal and N-terminal IDRs. The elicitin core domain was consistently below the disorder cut-off, indicating stable folding across all ELLs. Consistent with previous reports^39^, most *Albugo* ELLs were predicted to carry a glycosylphosphatidylinositol (GPI) membrane anchor (Figure 2D and E). Overall, our analysis indicates that IDRs are a predominant feature of *Albugo* ELLs, prompting us to investigate their functional role in this unique class of oomycete effectors.

### Intrinsic disorder in *Albugo* elicitin-like effectors enables the formation of amyloids

As microbe-host interactions are influenced by filamentous protein assemblies and intrinsic disorder is associated with amyloid formation^73^, we investigated whether IDRs in *A. candida* and *A. laibachii* ELLs drive the formation of amyloid fibrils. Amyloid formation was first predicted computationally, revealing a moderate correlation between intrinsic disorder scores and amyloidogenic potential (R^2^ = 0.356, p = 0.001; <u>Supp. Figure 15</u>). ELLs with higher disorder scores were generally predicted to be more amyloidogenic. To validate the impact of IDR conformational flexibility on amyloid fibrillation, we expressed all ELLs in an *Escherichia coli*-based system, the curli-dependent amyloid generator (C-DAG). It is designed to demonstrate fibril forming propensity of amyloidogenic target candidate proteins via color-phenotype-assays on plates containing Congo red, an amyloid specific dye. A pale colony phenotype indicates lack of amyloid formation, whereas a red colony phenotype reflects Congo red binding and amyloid formation. We observed a strong concordance between disorder prediction and colony phenotype, with few exceptions (Figure 3A and B). E1409 showed a positive phenotype despite low disorder, whereas E202 showed a negative phenotype despite high disorder, indicating that disorder propensity alone is not sufficient to predict amyloid formation.

**Figure 3:**
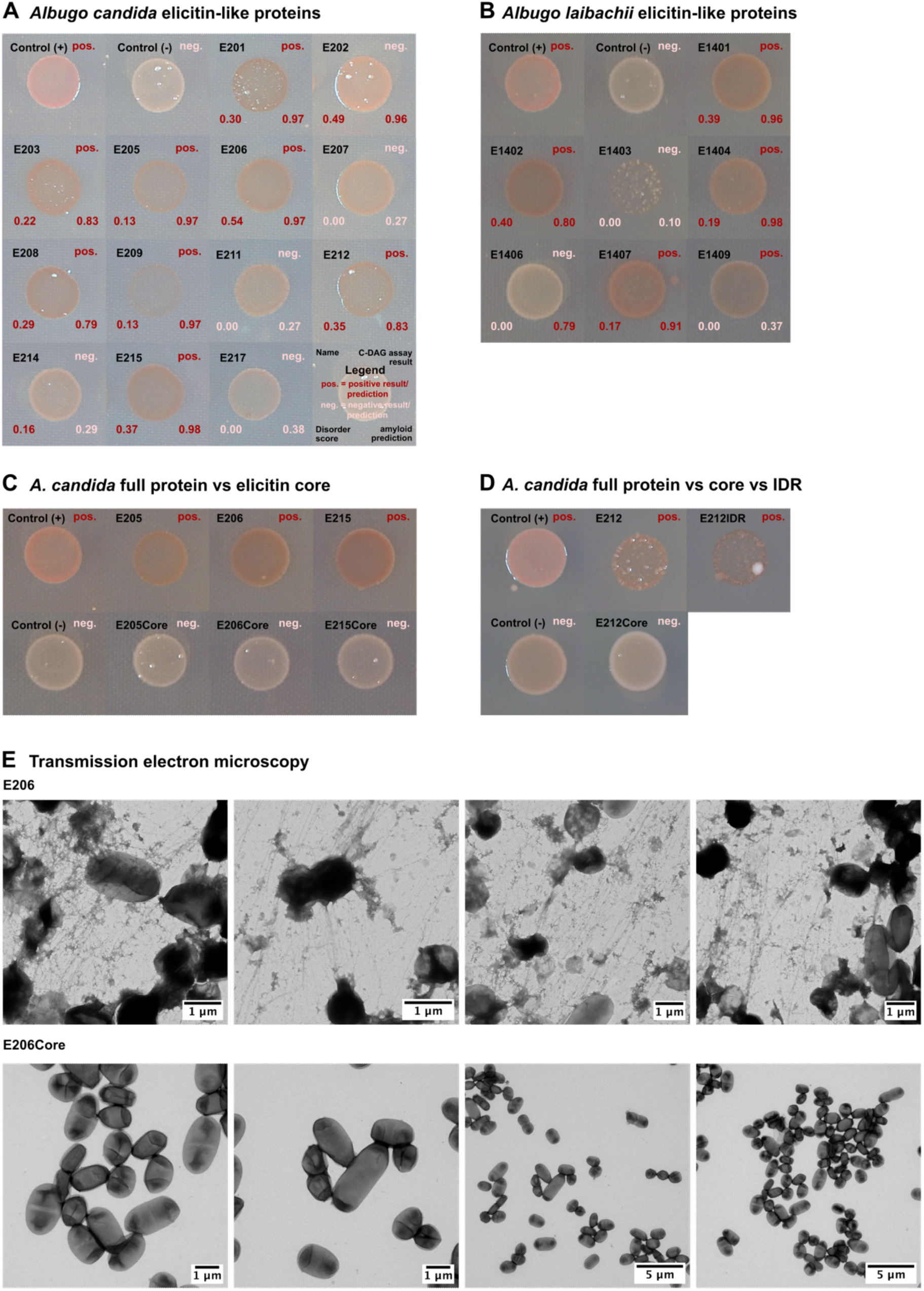
Analysis of amyloid-forming potential of elicitin-like proteins from *Albugo candida* (Nc2) and *Albugo laibachii* (Nc14). **A** Results of the curli-dependent amyloid generator (C-DAG) assay for elicitin-like protein (ELLs) from Nc2. Shown are pictures of the colony spots of *Escherichia coli* cells from the C-DAG system expressing ELL candidates on inducing Congo red agar. Positive (+) control is SUP35 an amyloidogenic prion from *Saccharomyces cerevisiae*, negative (-) control is the same protein without the aggregating domain. Protein name can be found in the top left corner, assay result in the top right, total disorder score in the bottom left and amyloid prediction score in the bottom right corner. Dark red indicates positive result/prediction, light red negative. Proteins are considered to contain intrinsically disordered regions (IDRs) when the disorder score is higher than 0. The amyloid prediction is considered positive when higher than 0.5 as suggested by the developer. **B** C-DAG assay results for ELL constructs from Nc14. **C** C-DAG assay results comparing full-length ELLs from Nc2 with their respective ordered core domains (Core), corresponding to the blue regions in Figure 2E. **D** C-DAG assay results from E212, core (Core) and disordered domain (IDR). In this assay a lower concentration of L-Arabinose (0.05%) was used in the plates to induce protein expression. **E** Transmission electron microscopy (TEM) images of *E. coli* cells from the C-DAG system expressing E206 and E206Core. Scale is indicated as a bar (1 µm image 1-6; 5 µm image 7-8). Images of positive (SUP35) and negative (SUP35 derivative) controls can be found in <u>Supp. Figure 16</u>.

To test whether highly disordered domains drive fibril formation, we generated derivatives of selected ELLs based on disorder levels. For E205, E206 and E215, selected based on strong amyloid phenotypes, we analyzed constructs containing only the elicitin core domain (ELI-1-like), excluding the N- and C-terminal IDRs. For E212, we generated constructs containing either only the C-terminal IDR (E212IDR) or the N-terminal core domain (E212Core). When comparing the phenotypes of the core constructs with those of their respective full-length ELLs, we found that the truncated ELLs lacking the IDRs had lost the ability to form amyloids, demonstrating that IDRs are required for amyloid formation in these ELLs (Figure 3C). Cells expressing E212IDR displayed impaired growth, likely due to the strong amyloidogenic nature of the protein overwhelming the cellular machinery. When E212IDR was expressed at low levels, growth improved and a red phenotype was observed, indicating that the IDR alone can assemble into amyloid fibrils (Figure 3D). We further examined fibril structures and validated C-DAG phenotypes using transmission electron microscopy (TEM). For this, we used E206 and E206Core due to robust growth, clear C-DAG phenotypes, and detection in the apoplastic proteomics. We observed abundant fibril formation, with long straight fibrils and mesh-like fibril structures in cells expressing full-length ELL E206, whereas no surface modifications or fibrils were detected in E206Core-expressing cells (Figure 3E, <u>Supp. Figure 16</u>). These results establish a structural link between intrinsic disorder and amyloid formation in *Albugo* ELLs.

### Intrinsic disorder shields elicitin core domains from recognition by plant receptors

To investigate whether intrinsic disorder in *Albugo* ELLs is relevant for pathogen-host interactions, we conducted infiltration assays in *A. thaliana* using selected ELLs to assess the influence of IDRs on PAMP properties. If IDRs function as molecular shields, they would be expected to reduce recognition of ELLs by plant PRRs and consequently attenuate cell death. We used the PURExpress^®^ cell-free system to express E212 and its derivatives E212Core and E212IDR, as well as E206 and E206Core. Lastly, E207 was included as a candidate with natively low disorder (Figure 2D). AlphaFold accurately predicted the structure of the ordered elicitin core domain, whereas IDRs appeared largely unstructured and exhibited low prediction scores (Figure 4A). SDS-PAGE analysis confirmed expression of E207, E206Core and E212Core as monomers of the expected size. Consistent with their amyloidogenic properties, E206, E212 and E212IDR additionally formed aggregates upon expression (<u>Supp. Figure 17</u>). Targeted LC-MS/MS analysis of peptides extracted from the aggregate band of E212 identified two peptides with 99% confidence and one peptide with 89% confidence. BLASTP homology search yielded a single hit for each peptide corresponding to E212 (<u>Supp. Figure 18</u>).

**Figure 4:**
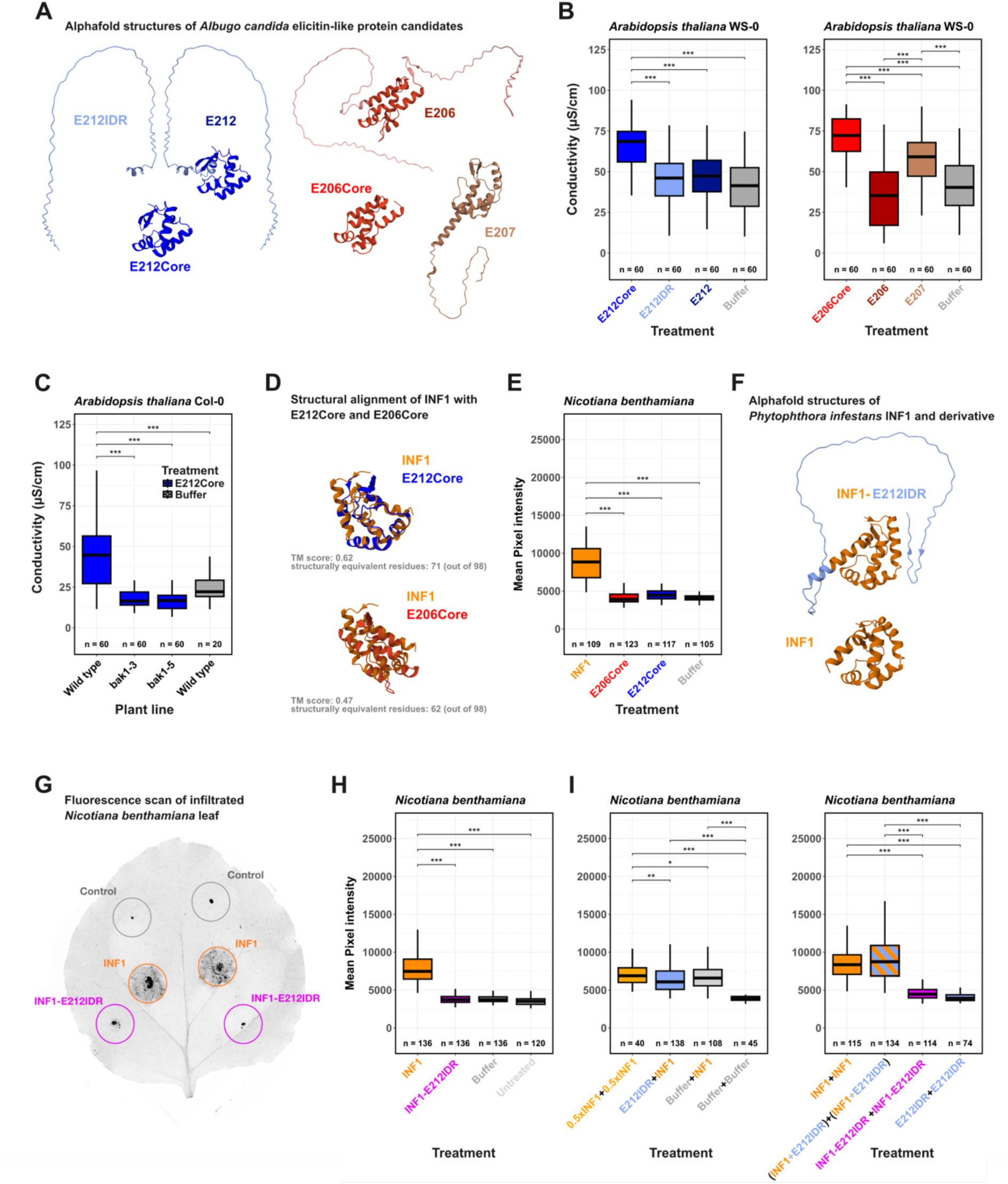
Cell death assessment after infiltration of elicitin-like candidates into Arabidopsis thaliana *and* Nicotiana benthamiana *leaves*. **A** Predicted AlphaFold 3 structures of elicitin-like (ELL) protein candidates and derivatives from *Albugo candida* Nc2. Colors represent different domains (matching the plots in subsequent figures). **B** Comparison of conductivity (as a proxy for cell death) of *A. thaliana* leaf areas infiltrated with ELL candidates and derivatives. Shown is the exemplary time point of three hours after measurement start, combining all three rounds (biological replicates). Colors indicate the treatments. The full-time courses for each round can be found in <u>Supp. Figure 19</u> and <u>Supp. Figure 20</u>. **C** Conductivity comparison of E212Core (blue) or buffer (grey) infiltrated Columbia-0 (wild type) and bak1 mutant lines, bak1-3 (SALK_034523) and bak1-5. Full-time courses in <u>Supp. Figure 21</u>. **D** Structural alignment of AlphaFold predicted structures of *Phytophthora infestans* INF1 and elicitin core domains from *A. candida* ELL candidates. RCSB structural alignment tool was used. Template modeling scores (TM-scores) are shown below the structures; TM >0.5 generally indicates the same protein fold. **E** Pixel intensity (as proxy for cell death) comparison of infiltrated *N. benthamiana* leaf areas imaged with a fluorescence scanner. *A. candida* elicitin core domains and INF1 were used for infiltration. **F** Predicted AlphaFold 3 structures of *P. infestans* INF1 and hybrid INF1 fused to the IDR of E212 from *A. candida* (E212IDR). **G** Exemplary *N. benthamiana* leaf infiltrated with INF1, INF1-E212IDR and buffer after fluorescence scanning. Colored circles indicate the infiltrated area, with the color showing the different treatments. **H** Pixel intensity comparison of *N. benthamiana* leaf areas infiltrated with INF1 and derivative. Experimental schematic is shown in <u>Supp. Figure 1</u>. **I** Pixel intensity comparison after sequential infiltrations with two different proteins (First infiltration (day 1) + Second infiltration (day 2)). (INF1+E212IDR) consists of a mixture of unfused INF1 and E212IDR. Experimental schematic can be found in <u>Supp. Figure 1</u>. **Statistical analysis:** lmer function was used to fit a linear mixed-effects model to the data, estimating the fixed effect of the treatment on the conductivity/pixel intensity while accounting for the random effect of the three biologically separate rounds. P-values were obtained from pairwise comparisons with the Tukey method using the emmeans function. Not significant comparisons are not displayed. *p < 0.05; **p < 0.01; ***p < 0.001.

The expressed *A. candida* ELL proteins, together with an expression reagents-only control, were used for infiltration assays in *A. thaliana*. Six days after infiltration, electrolyte leakage was measured via conductivity as a proxy for cell death and immune recognition. E212 and E212IDR, did not show significant difference in conductivity compared to the buffer control. In contrast, E212Core showed a significant increase in conductivity relative to both control and full-length protein (p < 0.0001). Similar results were observed for E206: full-length E206 resembled the control, whereas E206Core induced significantly higher conductivity (p < 0.0001). Increased conductivity was also observed for E207, a natively ordered ELL (Figure 4B, <u>Supp. Figure 19</u>, <u>Supp. Figure 20</u>). These results indicate that IDRs reduce recognition of elicitin core domains, consistent with a shielding function that attenuates plant defense responses.

To further characterize this molecular interaction, we performed infiltration assays in *A. thaliana* bak-1 mutants to test whether recognition of *A. candida* elicitin core domains is BAK1-dependent. In *S. microdontum* and *N. benthamiana,* elicitin recognition is mediated by BAK1-dependent PRRs^46,47^. Therefore, we infiltrated the bak1 mutants (bak1-3 and bak1-5) with E212Core and found loss of cell death induction (p < 0.0001), indicating recognition via a PRR that requires BAK-1 for initiating defense (Figure 4C, <u>Supp. Figure 21</u>). Despite their structural similarity to INF1, E212Core and E206Core did not induce cell death upon infiltration in *N. benthamiana,* in contrast to INF1 as measured through fluorescence scanning of the infiltrated areas (Figure 4D & E). These results suggest host-specific recognition of elicitin core domains and divergence in receptor systems.

Given the apparent specificity of core recognition, we next tested whether IDR-mediated shielding represents a transferable mechanism. For this, we fused E212IDR to INF1, the well-characterized ELI-1 known to cause cell death in *N. benthamiana* through recognition by the REL receptor (Figure 4F). Both INF1 and hybrid INF1-E212IDR were expressed via the PURExpress^®^ system. We additionally created a version with N- or C-terminal HiBit tags and verified their expression via the NanoGlo HiBit blotting system, showing monomers for INF1 and monomers as well as aggregates for INF1-E212IDR (<u>Supp. Figure 22</u>). We infiltrated INF1 and INF1-E212IDR into *N. benthamiana* and quantified cell death through fluorescence scanning of the infiltrated areas. As expected, native INF1 triggered cell death evidenced by an increased mean pixel intensity. In contrast, INF1-E212IDR did not induce cell death and resembled the buffer control (p < 0.0001), indicating that the E212 IDR prevents recognition of INF1 by REL (Figure 4G & H). Pre-infiltration or co-infiltration of E212IDR with INF1 did not prevent recognition, indicating that effective shielding requires covalent attachment of the IDR to the core protein (Figure 4I). The hybrid INF1-E212IDR exhibited a disorder profile similar to the native *P. infestans* elicitin INF2a (<u>Supp. Figure 12</u>), for which removal of disordered C-terminal regions restores recognition and induces cell death^74^. These results indicate that IDRs can conceal PAMP-associated recognition sites independent of the underlying core domain, enabling evasion of host immune responses. IDRs in ELLs therefore provide a selective advantage through molecular shielding, revealing a mechanism that contributes to the arms race between biotrophic pathogens and their host plants.

## Discussion

The coevolution of plant pathogens and their host constitutes an intricate molecular arms race whose depth and complexity have been studied for decades^75–78^. Understanding the underlying mechanisms is important for protecting economically relevant crops from infection^4^. In bacterial effectors, intrinsic disorder has emerged as a protein feature that can enhance fitness and promote host manipulation and immune evasion^15^. In this work, we therefore investigate whether similar principles operate in the secretome of plant pathogenic oomycetes and, particularly in their elicitin-like effectors^79–82^.

Functional characterization of IDRs in oomycete apoplastic effectors is still rare despite the high prevalence of IDRs in oomycete secretomes, as shown here, and similar abundance previously reported for bacterial and fungal apoplastic effectors^23,83^. In this study, we found that within the oomycete secretomes, elicitins are particularly enriched in IDRs, followed by RxLR effectors, for which disorder has already been linked to advantages in translocation^84^. In elicitins and ELLs, however, IDRs have received little attention, partly due to a focus on classical ELI-1 proteins such as INF1, which consist solely of the structured elicitin core and lack IDRs.

Remarkably, our analysis revealed that obligate biotrophic oomycetes are predicted to express a small number of elicitins, most of which are associated with intrinsic disorder and none of which are classified as ELI-1. This is consistent with their reliance on a living host, as intrinsic disorder may provide adaptive advantages for modulating host immunity. Classical ELI-1 elicitins, which are readily recognized by plant receptors, may therefore be selected against in obligate biotrophs. The reduced number of elicitins in these species may relate to their more compact proteomes and metabolic network^28,85^, while the enrichment in disorder may contribute to limiting host detection. Hemibiotrophs, by contrast, produce a wider variety of elicitins, potentially benefiting from reduced recognition during early biotrophic stages while being overall less dependent on sustained host viability. In *Phytophthora* spp., ELLs are often expressed during early infection stages such as in zoospores, including proteins with disorder profiles similar to *A. candida* E212 and E206 (<u>Supp. Figure 11</u>). In contrast, highly recognizable ELI-1 proteins such as INF1 are predominantly expressed during later mycelial stages^51,86^. Together, these observations suggest that intrinsic disorder in elicitins and ELLs may be a characteristic feature of the biotrophic phase, with distinct elicitin types deployed at different infection stages depending on the pathogen’s requirements and environmental conditions such as sterol availability^41^. Considering the dependence of sterol auxotrophs on elicitin secretion, a more detailed analysis of elicitin expression patterns and the diversity of IDRs across oomycete species will provide important insights into how pathogens balance nutrient acquisition with avoidance of host immune recognition during infection.

The obligate biotrophs *A. candida* and *A. laibachii* predominantly secrete ELLs containing IDRs. To investigate the functional relevance of these IDRs, we first examined a structural property commonly associated with intrinsically disordered proteins, namely the ability to form amyloid fibrils. The concept of functional amyloids across all domains of life has gained increasing attention^87,88^, with emerging evidence for roles in host-microbe interactions^89^ including within the plant microbiome^20^. Amyloids are fibrillar protein assemblies that often originate from proteins containing stretches of disorder^17^. Using the C-DAG system, we screened all ELLs from *A. candida* and *A. laibachii* for amyloidogenic properties. The assay revealed a high prevalence of amyloid-forming capacity among ELLs from both *Albugo* species, which broadly correlated with their disorder content. For one candidate, we found filament formation despite low disorder, indicating that additional sequence features beyond overall disorder contribute to amyloid formation. It has been suggested that specific sequence motifs, once exposed through unfolding, can drive aggregation^90^. For candidates with high disorder, we found that the IDRs are required for fibril formation, as removal of these regions abolished amyloid formation. This suggests that intrinsic disorder facilitates amyloid assembly in ELLs by enabling exposure of aggregation-prone sequence motifs within these proteins. For fungal spores, surface amyloids have been shown to mask spores from host immune recognition^91,92^. In addition, the availability of compounds, such as cell wall components, can influence amyloid assembly^93^. In the context of elicitins, which contain conserved core domains prone to immune recognition, similar properties would be advantageous. Self-assembly into fibrils linked to availability of sterol could, in principle, reduce exposure of immunogenic regions during the biotrophic phase. There is also evidence that elicitins may be tethered to the cell surface or cell wall^94^. Hyphal surface amyloids are common and, at least in filamentous fungi, have been linked to pathogenicity^95^. Many ELLs carry predicted GPI anchor and O-glycosylation sites in their C-terminal regions^51^, consistent with the pattern observed in *Albugo* ELLs. This could enable the formation of surface-associated fibrils from elicitin monomers attached to the pathogen surface. By analogy to systems such as the bacterial toxin microcin^96^, such fibrils might act as a reservoir of elicitin molecules, potentially allowing regulated release depending on environmental conditions, for example sterol availability.

To examine how IDRs influence PAMP activity, we selected ELL candidates from *A. candida* for functional analysis. Elicitin expression and release are required for sterol acquisition^40^. In the context of the molecular arms race, plants have evolved pattern recognition receptors that detect the conserved elicitin core domain, thereby selecting for pathogen counter-strategies^1,46,47^. In line with this, *P. infestans* RxLR effectors such as AVR2 and AVRblb2 contain IDRs or IDR-associated motifs that have been linked to immune evasion and host-target manipulation^97^. While the classification of elicitins into virulence and avirulence factors remains ambiguous^39^, studies on AVR2 illustrate how IDRs can contribute to immune evasion. The virulent AVR2 variant contains disordered regions that promote evading detection by resistance proteins, whereas the avirulent form lacks disorder and interacts with the R2 protein^98^. Avrblb2 contains accessible motifs within IDRs that interact with a plant ion sensor and thereby interfere with plant defense signaling through host mimicry^99^.

In ELLs, our data support a model in which IDRs shield the conserved core domain from detection by pattern recognition receptors such as ELR and REL. Such receptors have been described in *S. microdontum*^46^ and *N. benthamiana*^47^, however, in *A. thaliana,* similar receptors have been proposed but not yet characterized. Our findings show that the conserved elicitin core domains from *A. candida* elicitins is recognized by *A. thaliana* in a BAK1-dependent manner, leading to cell death responses and suggesting the presence of a PRR that detects *Albugo*-specific elicitin core regions. However, recognition is abolished when the conserved core is shielded by IDRs, demonstrating that these additional domains are functionally relevant. While infiltration of *A. candida* elicitin core domains into *N. benthamiana* indicated that core recognition is host-specific, we found that the shielding function of IDRs is maintained across systems and is independent of specific core domains. INF1 fused to the IDR from E212 no longer induces cell death in *N. benthamiana,* indicating that the INF1 domain is no longer recognized by the pattern recognition receptor REL. Notably, shielding requires physical linkage, as infiltration of the IDR alone did not affect the cell death response. This is consistent with observations for INF2a, an ELI-2/3/4 protein with a conserved core and a C-terminal IDR. Native INF2a including its C-terminal IDR, induce only little cell death when expressed in *N. benthamiana*. However, removal of the C-terminal domain results in strong necrosis^74^.

Effector shielding represents a potential strategy in the molecular arms race and may also operate in other oomycete effectors. In our analysis, members of the cysteine-rich secretory proteins antigen 5 and pathogenesis-related 1 (CAP) superfamily were also strongly associated with IDRs. In plant-pathogen interactions, these apoplastic effectors remain understudied, however, CAP1 from *P. sojae* is secreted during early stages of infection and can trigger immune responses. Similar to INF1, CAP1 contains little intrinsic disorder and is recognized by a plant membrane-localized LRR-RLP associating with BAK1 and SORBIR1. The presence of multiple CAP domain-containing effectors with high disorder in *A. candida* raises the possibility that IDR-mediated shielding may also operate in this effector class.

Overall, these findings identify intrinsic disorder as a key feature of elicitin and ELL effectors that reduces immune recognition during the biotrophic phase, consistent with an IDR-mediated shielding mechanism, and thereby promotes successful infection. IDR-associated functions thus contribute to pathogen persistence, and understanding these mechanisms may support the development of effective strategies for plant protection. Our findings further support a cross-kingdom relevance of intrinsic disorder in apoplastic effectors and highlight the importance of considering the structure-function-continuum in effector biology.

## Supporting information

Compiled Supplemental Information

## Resource availability

### Lead Contact

Eric Kemen is managing the used resources and may be contacted for further questions at.

### Materials Availability

Materials and strains are available at the Microbial Interactions in Plant Ecosystems Lab at the University of Tuebingen. Lead contact may be contacted regarding inquiries.

### Data and Code Availability

Data was compiled and curated by Monja Schmid. All data is deposited in a zenodo repository under the following doi 10.5281/zenodo.17936530. This includes all experimental results discussed and used in the figures, the code for the bioinformatics analysis and creation of the plots. All data regarding apoplast proteomics is curated and deposited under the following doi http://doi.org/10.57754/FDAT.8k1w5-2yh42.

## Acknowledgement

We would like to acknowledge support from the European Research Council (ERC) under the DeCoCt research program (grant agreement: ERC-2018-COG 820124) and the TRR 356 PlantMicrobe. Metabolite analytics in the ZMBP central facilities unit was funded by the DFG (Project number 442641014). We would like to thank Moritz Urban and Farid El Kasmi for their assistance with the pixel intensity measurement, Libera Lo Presti for her valuable feedback and suggestions and Rebecca Stahl, Lorenz Henneberg and Sandra Richter from the microscopy unit of the ZMBP (funded by the DFG under the project number INST 37/900-1 FUGG).

## Author contributions

Conceptualization, M.S., D.G.-P., A.K. and E.K.; Methodology, Software and Formal Analysis, M.S., D.G.-P. and S.Q.; Investigation, M.S., and E.R.-L.; Validation, M.S., A.K. and E.K; Visualization, M.S. and S.Q.; Data Curation, M.S. and E.K.; Resources, E.K.; Writing - Original Draft, M.S.; Writing - Review and Editing, M.S., D.G.-P., S.Q., A.K., E.R.-L. and E.K.; Supervision, A.K. and E.K.; Project Administration and Funding Acquisition, E.K.

