## Supplementary material for "Intrinsic disorder in elicitin-like effectors: Molecular shields in the arms race of biotrophic pathogens": Compiled Supplemental Information

### Supplementary figures

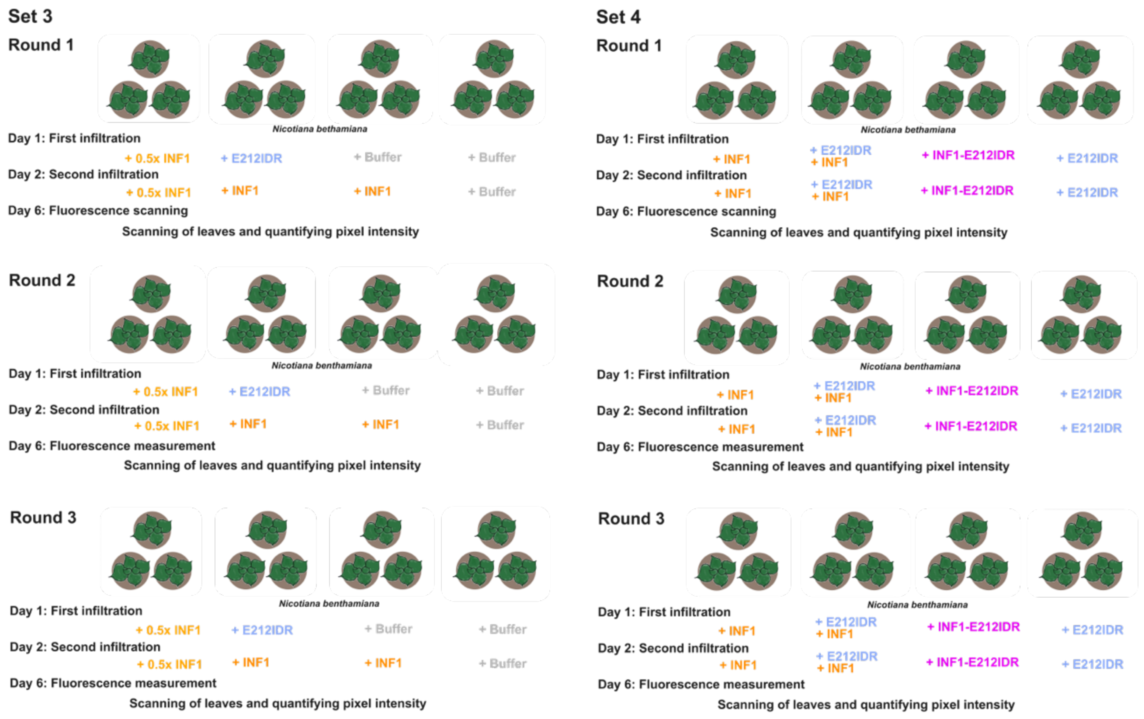

**Supp. Figure 1: Experimental overview of *Nicotiana benthamiana* infiltrations.**
Schematic of the experimental setup of *N. benthamiana* infiltrations Set 3 and 4 with elicitin-like protein candidates from *Albugo candida*, INF1 from *Phytophthora infestans* and hybrid INF1-E212IDR for cell death quantification. Addition of IDR to the protein name indicates only the disordered part was infiltrated and Core indicates only the elicitin core domain was used. Experiments are split into sets and rounds, which were conducted independently of each other.

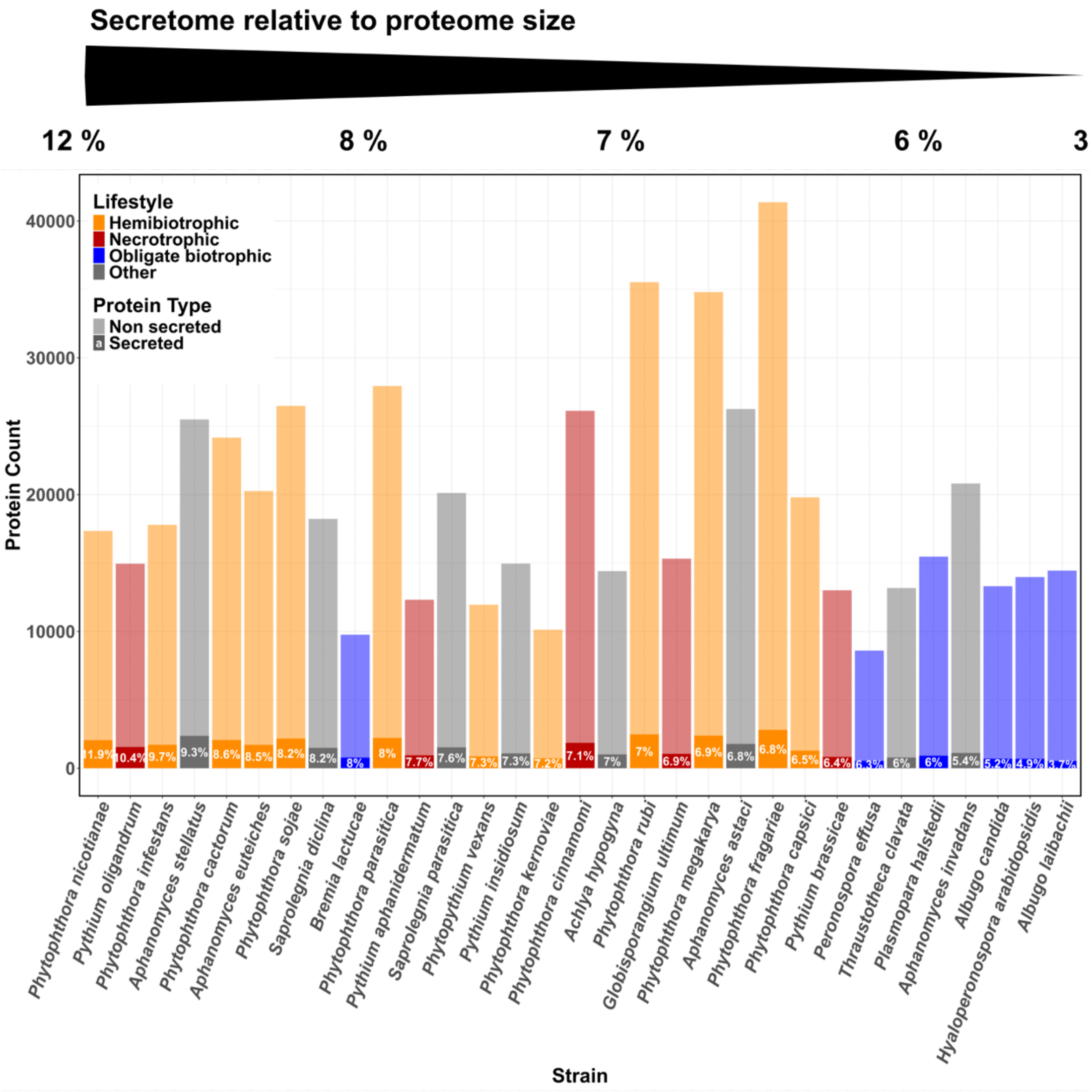

**Supp. Figure 2: Proteome and secretome size across oomycete strains.**
Dark shade indicates classical secretome as predicted with SignalP6.0, light shade indicates non secreted proteins; the number shows the exact percentage of secreted proteins relative to the overall proteome size per individual strain. Colors represent the lifestyle. The bars are sorted by percentage of the secretome from highest to lowest, as also indicated by the bar on top.

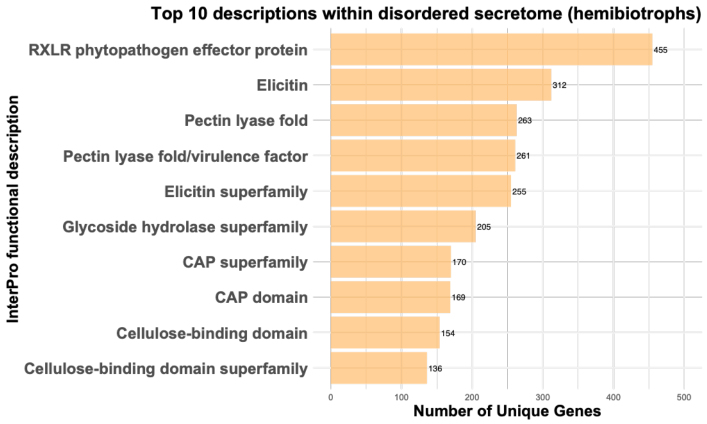

**Supp. Figure 3: Occurrence of functional descriptions within the disordered secretome of hemibiotrophs.**
Shown are the ten most frequent functional descriptions (InterProScan) within the disordered portion (disorder score >= 0.2) of the secretome in hemibiotrophs.

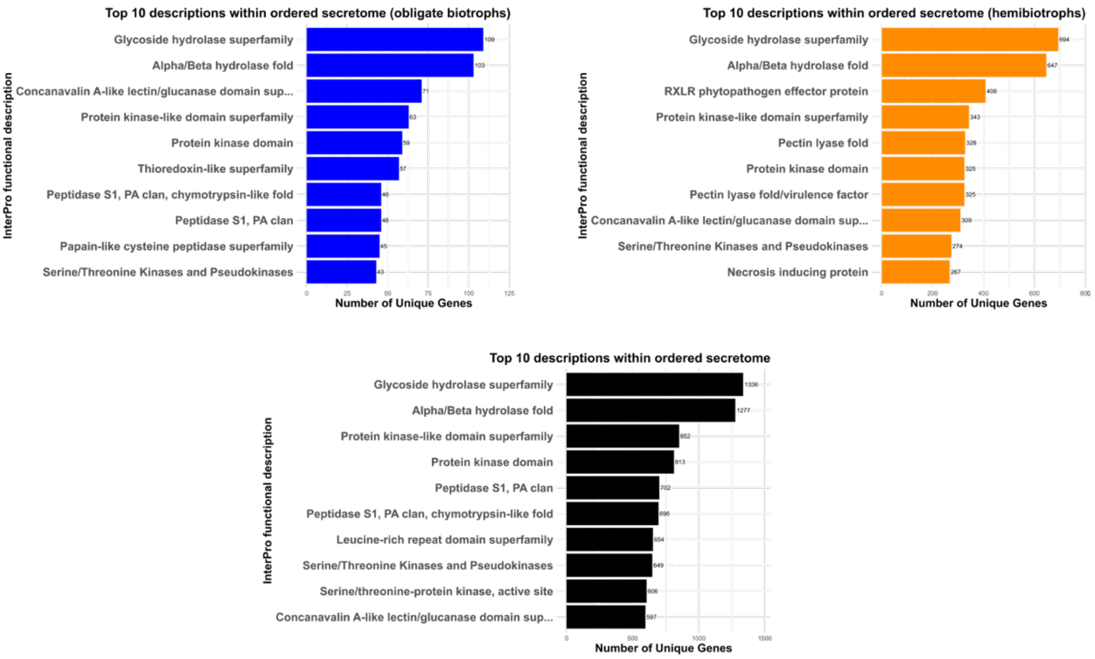

**Supp. Figure 4: Occurrence of functional descriptions within ordered secretomes.**
The black barred figure shows the ten most frequent functional descriptions (InterProScan) within the ordered portion of the secretome (disorder score < 0.2) across all lifestyles; the blue shows descriptions within the ordered portion across obligate biotrophs, and the orange shows the hemibiotrophic ordered descriptions.

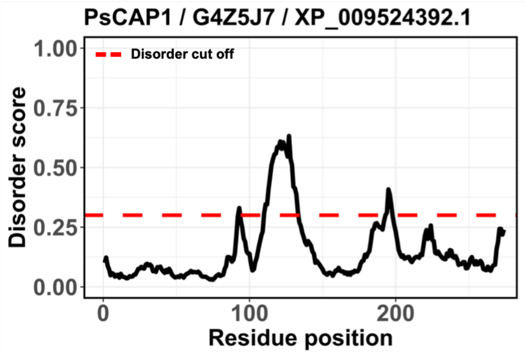

**Supp. Figure 5: Disorder profile of PsCAP1 from *Phytophthora sojae*.** The disorder cut off is shown as a red dashed line. Accession numbers can be found in the plot title.

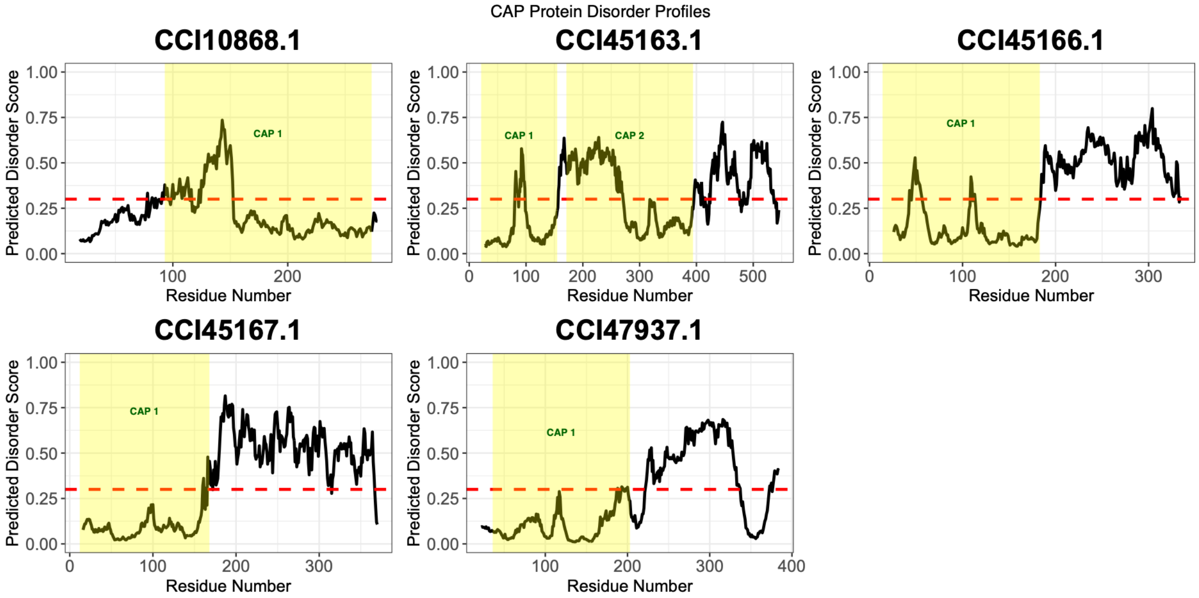

**Supp. Figure 6: Disorder profiles of proteins from *Albugo candida* annotated with a CAP domain.** The red dashed line shows the disorder cut off. The CAP domains are highlighted in yellow. The protein accession can be found in the individual plot titles.

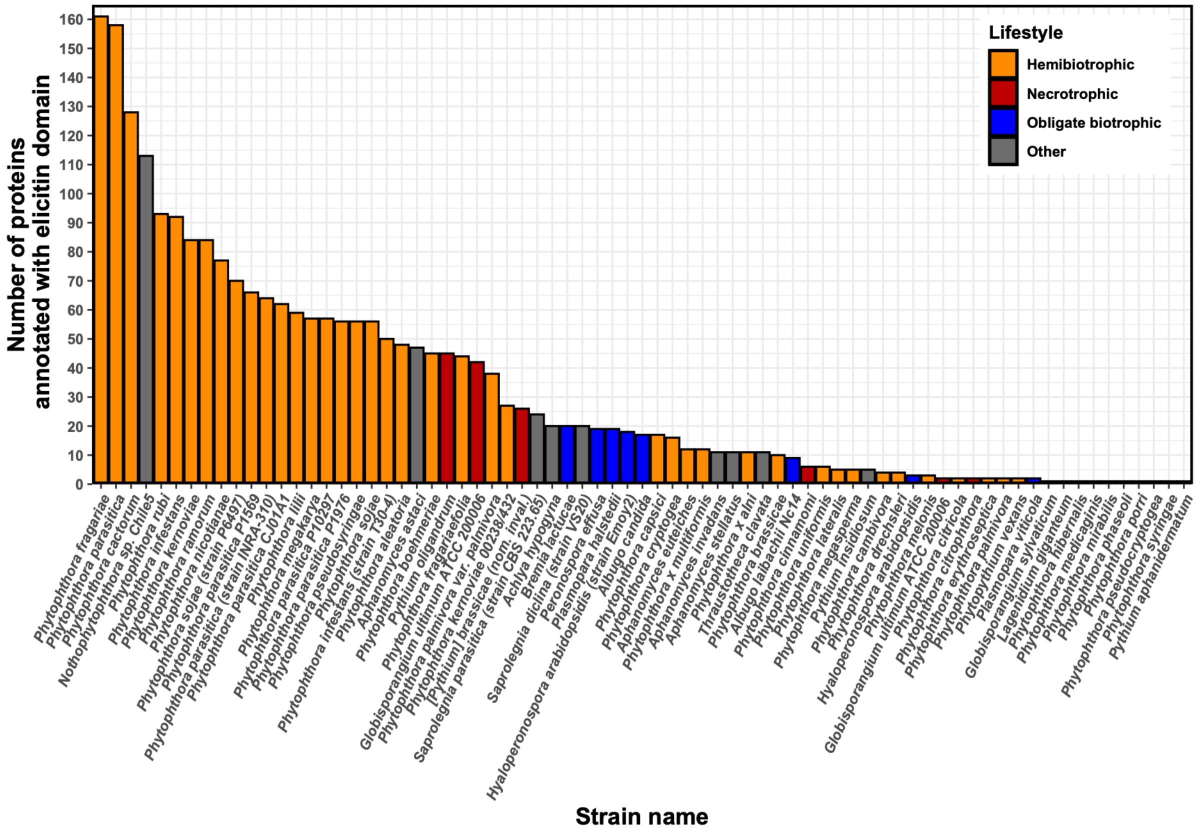

**Supp. Figure 7: Elicitin count per oomycete species.**
Bar plot showing the number of proteins annotated with an elicitin domain according to InterPro for each oomycete species. Colors represent lifestyle.

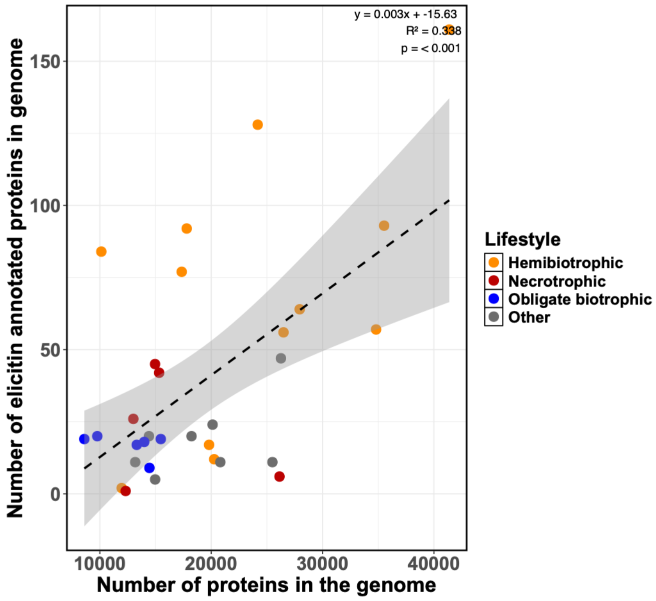

**Supp. Figure 8: Analysis of proteome size and elicitin count correlation.**
Correlation between the count of elicitins per strain and the count of proteins in the overall proteome of the respective strain. The dotted line shows the linear regression line following the formula y ~ x; the grey area shows the confidence interval around the regression line. Colors represent lifestyles.

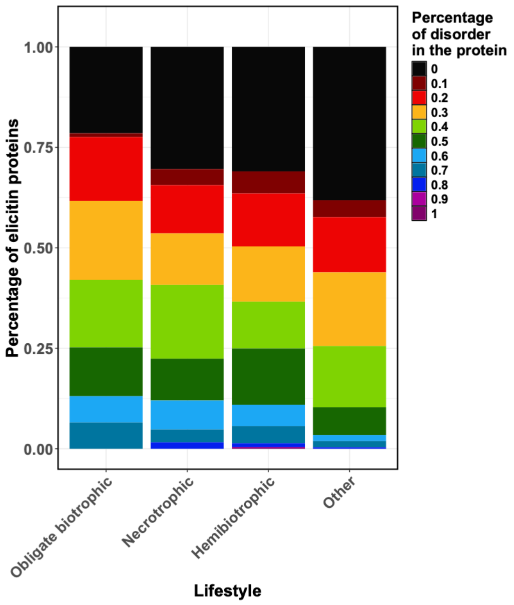

**Supp. Figure 9: Detailed disorder distribution within proteins annotated as elicitins.** Stacked bar plot showing the relative percentage of elicitin proteins with a certain level of disorder for each lifestyle. The colors show the disorder levels, ranking from no disorder at all (0, black) to complete disorder (1, purple).

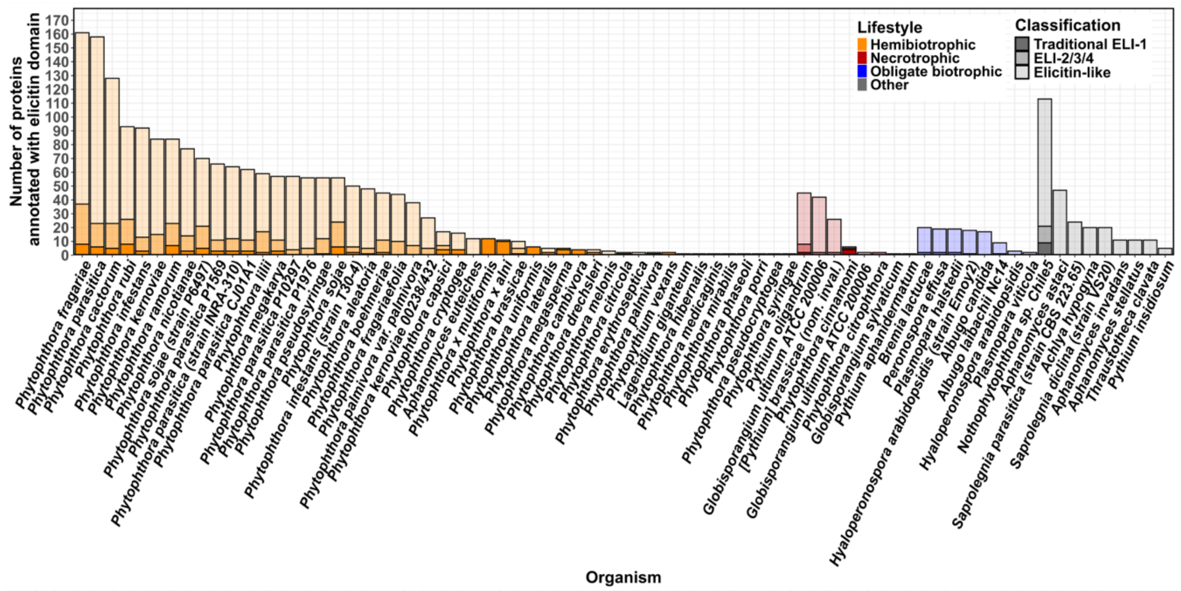

**Supp. Figure 10: Elicitin classification across all available strains.** Classification of proteins with annotated elicitin domain per species considering all strains in the dataset. Traditional class 1 elicitins (ELI-1) are defined as having a conserved core domain (96-100 amino acids with six cysteine residues) and no additional domains (dark shade). Class 2/3/4 elicitins (ELI-2/3/4) have the same core domain but exhibit additional domains (mid-tone shade). Elicitin-like proteins (ELLs) have more variable core and appending domains (light shade). Colors represent the different oomycete lifestyles.

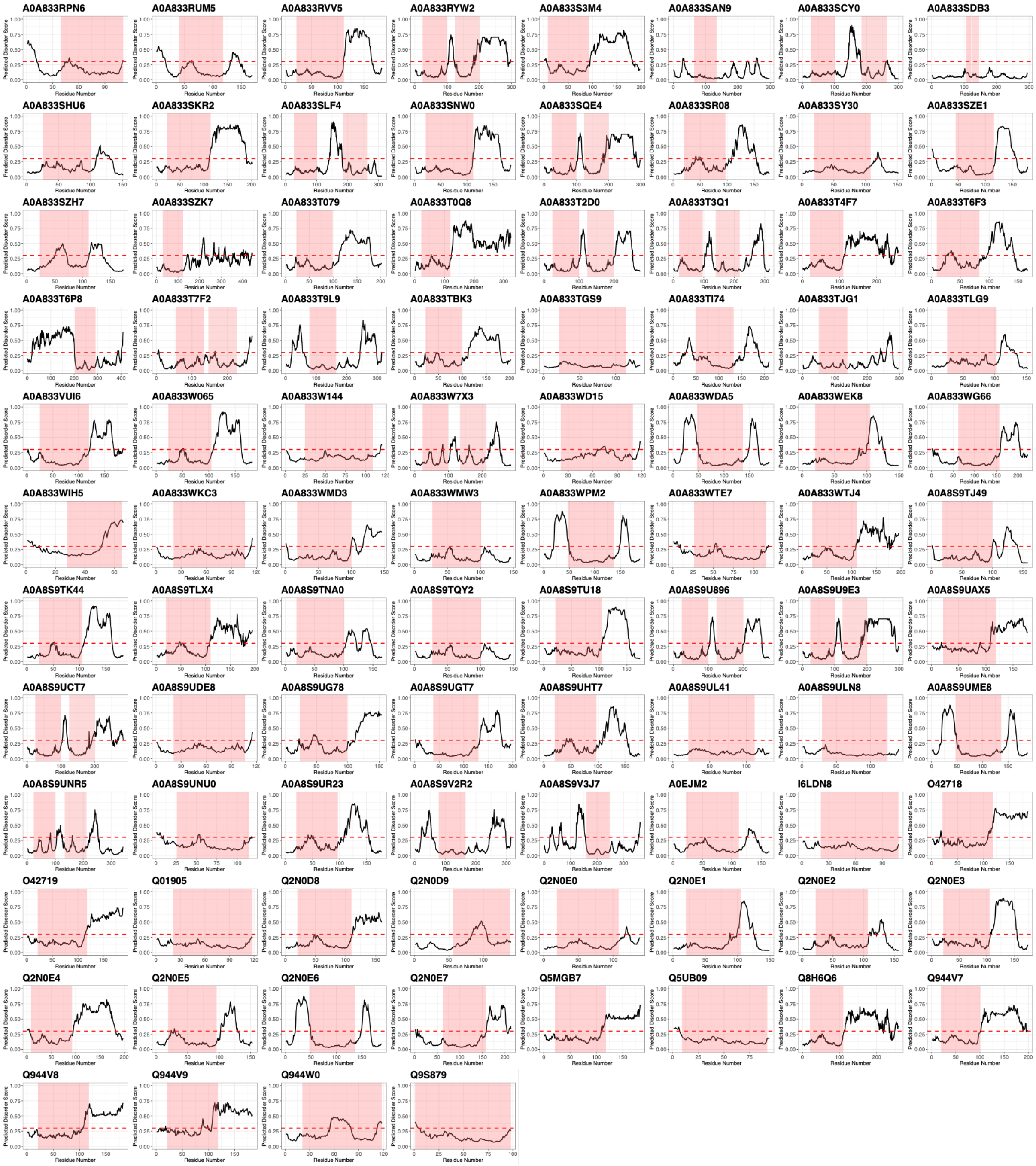

**Supp. Figure 11: Disorder profiles of *Phytophthora infestans* proteins annotated as elicitins.** Protein accessions are shown in the individual plot titles. The red highlighted area shows the annotated elicitin core domain. The red dashed line shows the disorder cut off. Well studied proteins include: INF1 (Q01905), INF2A (O42718), INF2B (O42719), INF4 (Q944W0), INF6 (Q5MGB7).

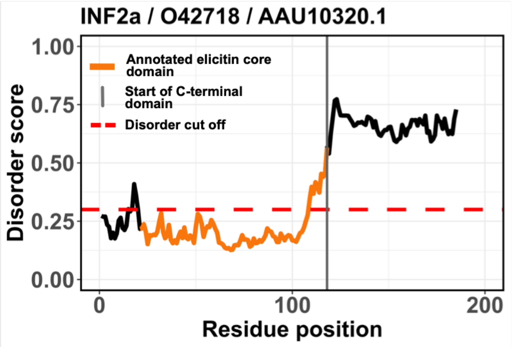

**Supp. Figure 12: Disorder profile of INF2a from *Phytophthora infestans*.** The annotated elicitin core domain is highlighted in orange, the start of the C-terminal domain is shown as a grey intersection line. The disorder cut off is shown as a red dashed line. Accession numbers can be found in the plot title.

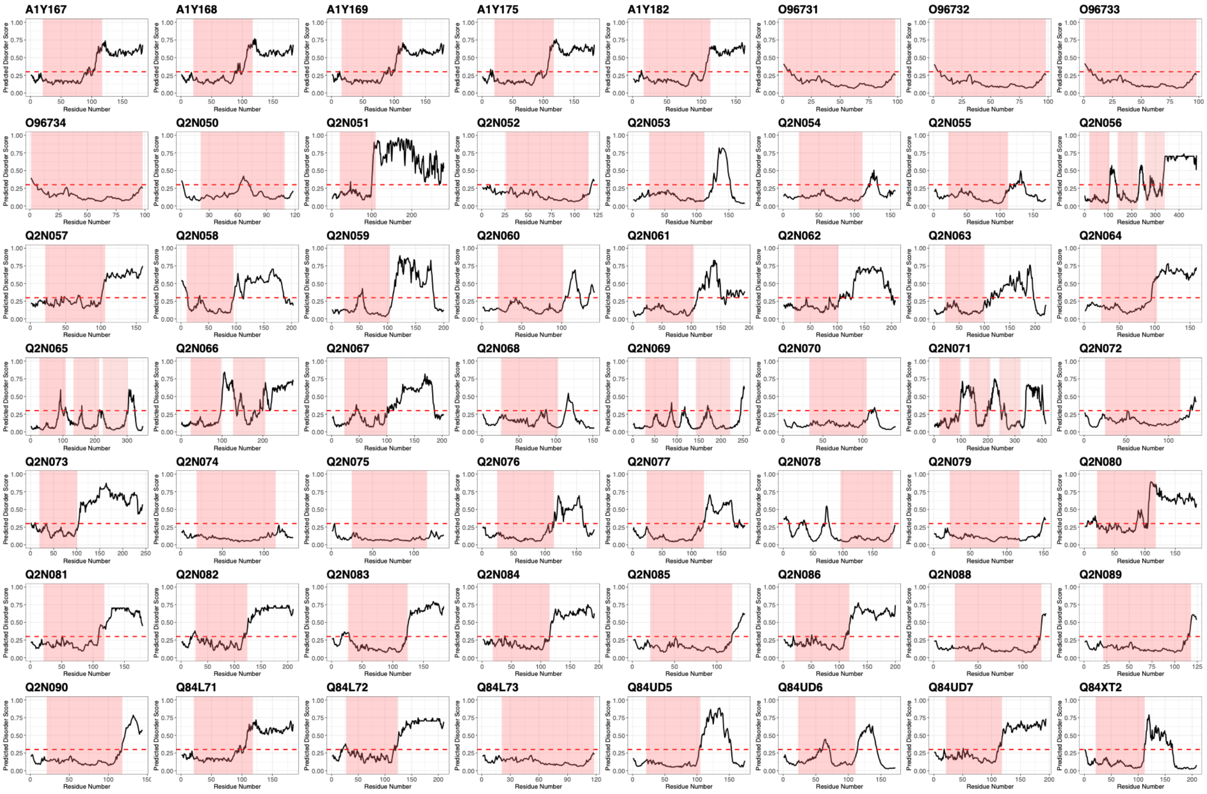

**Supp. Figure 13: Disorder profiles of *Phytophthora sojae* proteins annotated as elicitins.** Protein accessions are shown in the individual plot titles. The red highlighted area shows the annotated elicitin core domain. The red dashed line shows the disorder cut off. Well studied proteins include: SOJX (Q84UD6), SOJY (Q84UD5), SOJA (O96731, O96732, O96733, O96734), SOJB (Q84L73), SOJ2 (Q84UD7), SOJ3 (Q84L72), SOJ6 (Q84L71)[100].

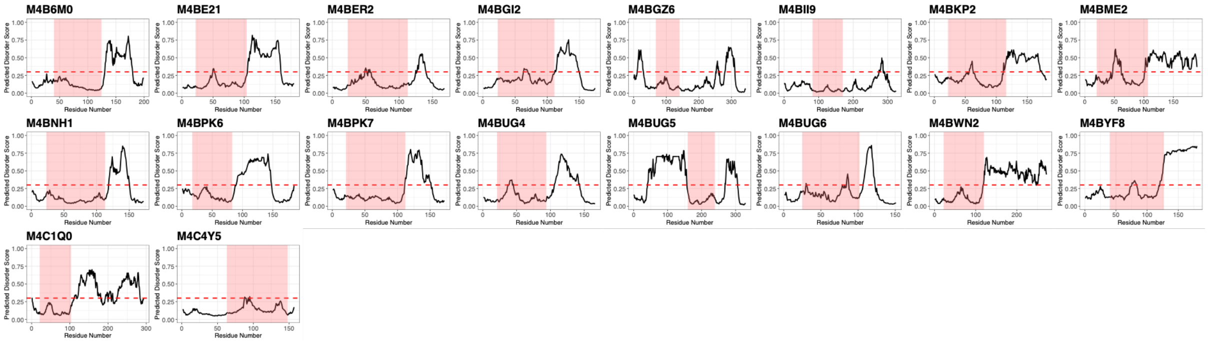

**Supp. Figure 14: Disorder profiles of *Hyaloperonospora arabidopsidis* Emoy2 proteins annotated as elicitins.** Protein accessions are shown in the individual plot titles. The red highlighted area shows the annotated elicitin core domain. The red dashed line shows the disorder cut off.

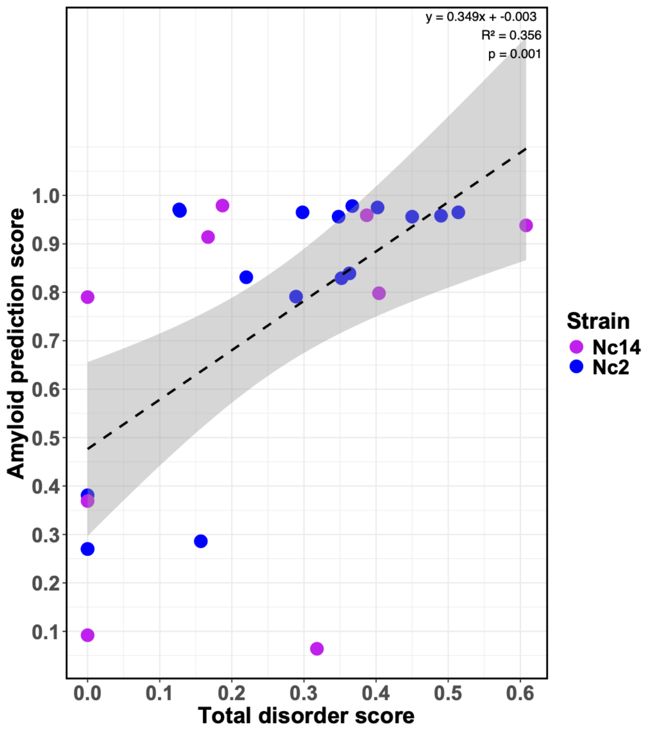

**Supp. Figure 15: Correlation between amyloid and disorder prediction scores for elicitin-like proteins from *Albugo candida* (Nc2) and *Albugo laibachii* (Nc14).** Correlation between the total disorder score and the amyloid prediction of the respective elicitin proteins. The dotted line shows the linear regression line following the formula y ~ x; the grey area shows the confidence interval around the regression line.

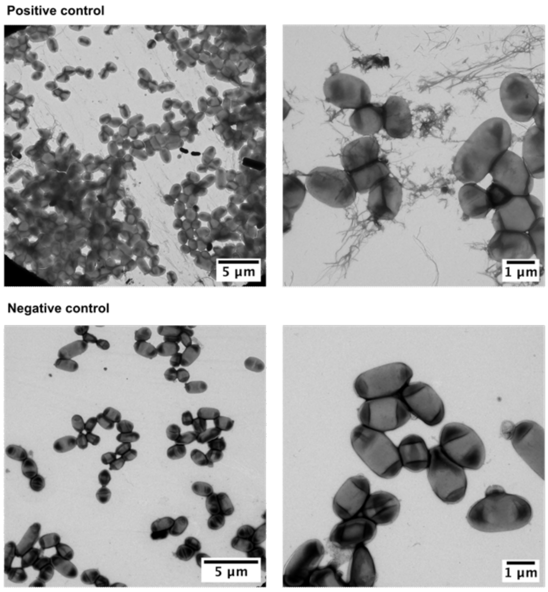

**Supp. Figure 16: Transmission electron microscopy images of controls.** Transmission electron microscopy (TEM) images of *Escherichia coli* cells from the C-DAG system expressing the positive control SUP35 on the top and the negative control SUP35 without aggregation domain on the bottom. Scale bar shows 5 µm in the images on the left and 1 µm in the images on the right.

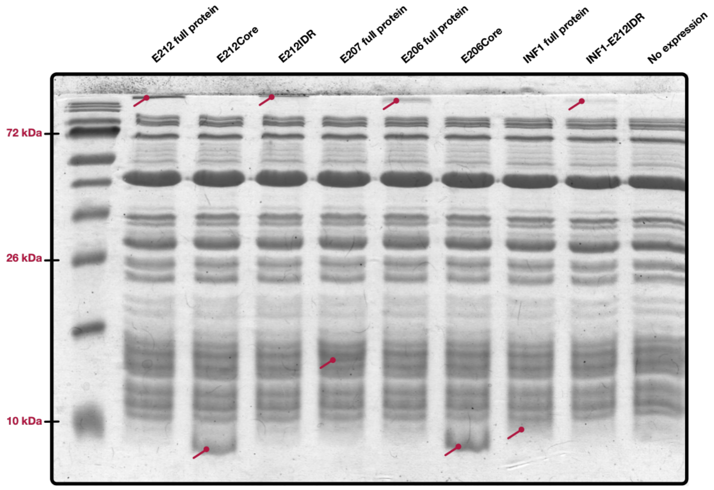

**Supp. Figure 17: SDS PAGE analysis showing elicitin expression.** SDS PAGE (18%) analysis of elicitin candidates expressed via the PURExpress® system. Red labels mark the relevant band. The last lane shows the control containing only the expression reagents but no expressed protein. The expected weights are as follows: E212: 19.61 kDa; E212Core: 9.74 kDa; E212IDR: 10.02 kDa; E207: 16.47 kDa; E206: 22.64 kDa; E206Core: 9.96 kDa; INF1: 10.46 kDa; INF1-E212IDR: 20.46 kDa.

Confidence: 89% 99% 99%

Sequence coverage: NYACDIPQIQGTLLPNGTEYNNICQSKSGYDIFSLDKYPNEEQVQVLSHTRECTDVLNQINSRANQLIQCDVNINGTNLSYGWLIS
TWLMGKTGNPSNETDSLTGENGSYEVEPSSADNSTSAIDSESKDEDVGKSKIQKKKEQKPAHASASTESRATSVCFSFVVGGF
FAFVVSIATV

**Supp. Figure 18: Sequence coverage of the elicitin candidate E212 expressed through PURExpress and extracted from the aggregation band after the SDS PAGE.**
Targeted LC–MS/MS analysis of the proteolytic digest of the E212 protein, expressed via PURExpress system and extracted from aggregation band after SDS PAGE, identified two peptides with 99% confidence (green) and one peptide with 89% confidence (orange), corresponding to an overall sequence coverage of 20.1%. All MS/MS spectra were manually inspected to confirm peptide authenticity. A BLASTP homology search of the high-confidence peptide region (26 amino acids; shown in green) against the NCBI NR protein database yielded a single hit with 100% identity to the hypothetical protein ABG067_001974 from *Albugo candida*, confirming sequence specificity.

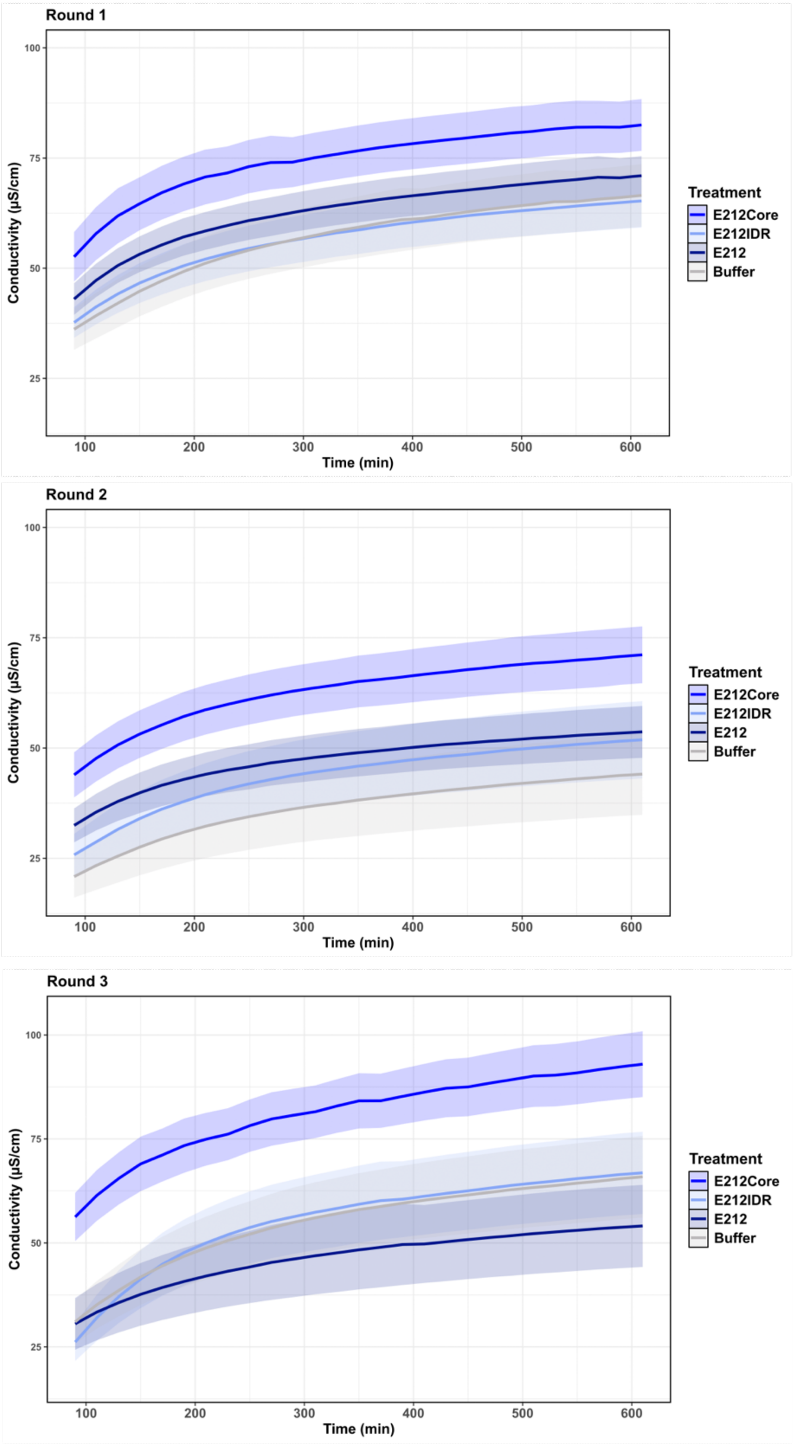

**Supp. Figure 19: Detailed time course on conductivity measurement with E212 constructs.** Conductivity measurement over time of *Arabidopsis thaliana* leaf areas infiltrated with constructs of E212. The rounds show the independent experimental set ups. The colors indicate the different treatments. Lines show the mean conductivity for each treatment across replicates at each timepoint; the shaded ribbons represent the 95% confidence interval (CI) around the mean (t-based CI calculated from the sample standard deviation and number of observations at each timepoint).

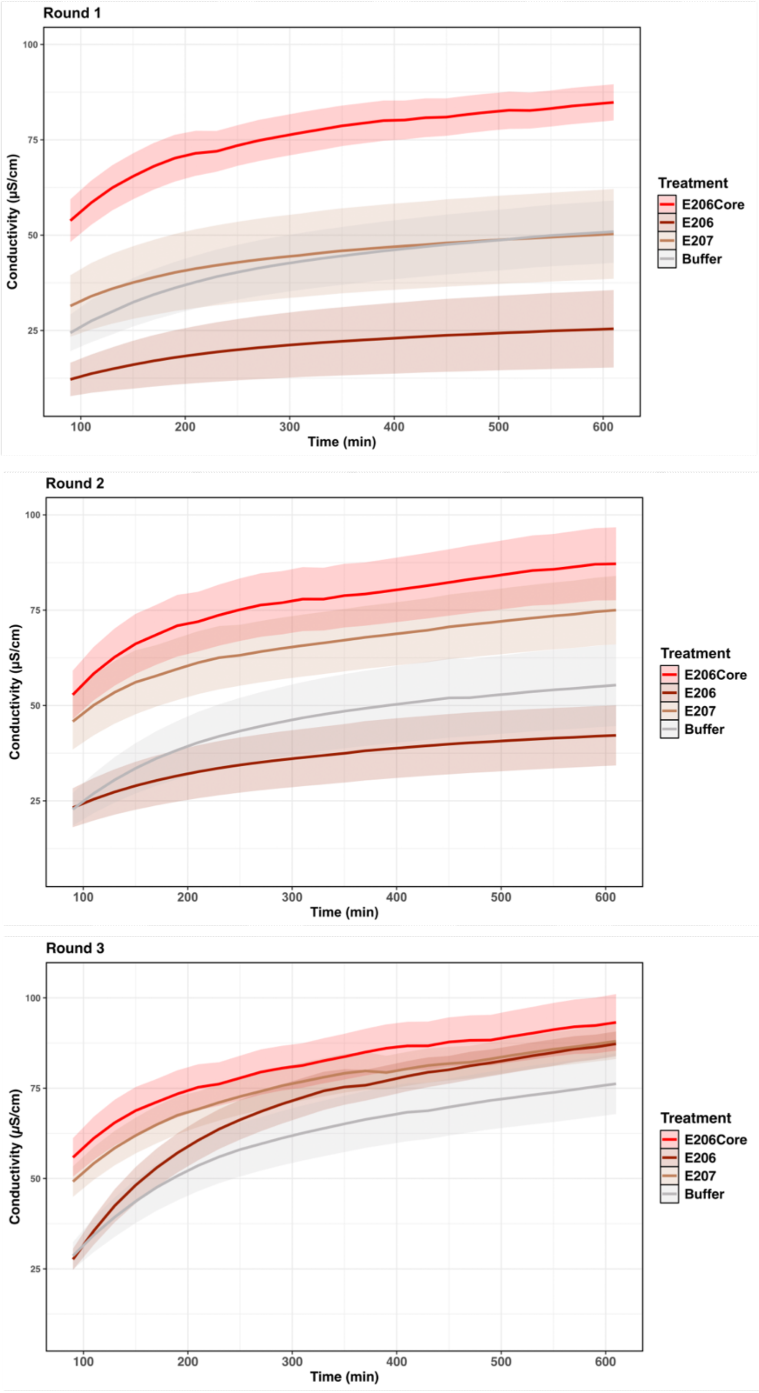

**Supp. Figure 20: Detailed time course on conductivity measurement with E206 constructs and E207.** Conductivity measurement over time of *Arabidopsis thaliana* leaf areas infiltrated with constructs of E206 and E207. The rounds show the independent experimental set ups. The colors indicate the different treatments. Lines show the mean conductivity for each treatment across replicates at each timepoint; the shaded ribbons represent the 95% confidence interval (CI) around the mean (t-based CI calculated from the sample standard deviation and number of observations at each timepoint).

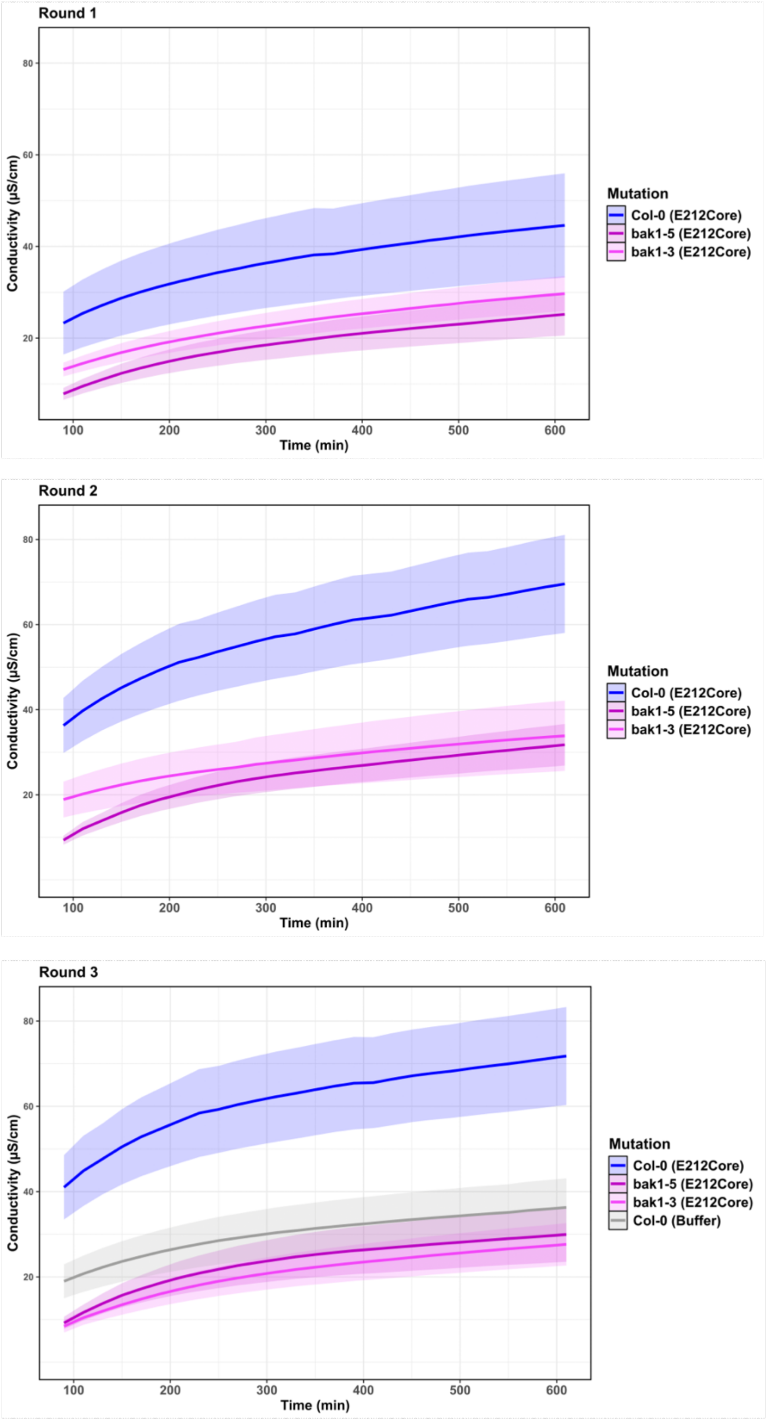

**Supp. Figure 21: Detailed time course on conductivity measurement with bak1 mutants.** Conductivity measurement over time of *Arabidopsis thaliana* leaf areas infiltrated with constructs of E212Core and buffer. The rounds show the independent experimental set ups. The colors indicate the different plant lines (Col-0 wildtype and bak1 mutant lines). Lines show the mean conductivity for each treatment/plant line across replicates at each timepoint; the shaded ribbons represent the 95% confidence interval (CI) around the mean (t-based CI calculated from the sample standard deviation and number of observations at each timepoint).

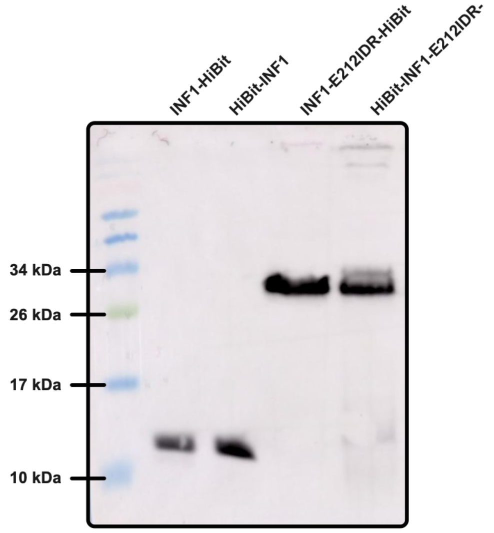

**Supp. Figure 22: Western blot analysis of successful expression of tagged INF1 and INF1-E212IDR.**

Developed membrane showing western blot result after SDS PAGE (18%) of proteins expressed via the PURExpress system. Proteins are tagged with Promega HiBit tag; the NanoGlo Blotting system was used to visualize the proteins on the membrane. Expected size: INF1-HiBit: 11.8 kDa; INF1-E212IDR-HiBit: 21.8 kDa.

#### Supplementary tables

**Supp. Table 1: Proteome overview of oomycete species.** The table contains the counts of overall proteins in the proteome, counts of how many proteins are predicted to be classically secreted (SignalP6.0), and the count of proteins annotated with an elicitin domain per strain.

| Strain | Elicitin count | Lifestyle | Proteome count | Secretome count | Secretome percentage |
|---|---|---|---|---|---|
| *Achlya hypogyna* | 20 | Other | 14406 | 1015 | 7.05 |
| *Albugo candida* | 17 | Obligate biotrophic | 13310 | 695 | 5.22 |
| *Albugo laibachii* | 9 | Obligate biotrophic | 14449 | 540 | 3.74 |
| *Aphanomyces astaci* | 47 | Other | 26259 | 1783 | 6.79 |
| *Aphanomyces euteiches* | 12 | Hemibiotrophic | 20265 | 1720 | 8.49 |
| *Aphanomyces invadans* | 11 | Other | 20816 | 1119 | 5.38 |
| *Aphanomyces stellatus* | 11 | Other | 25494 | 2374 | 9.31 |
| *Bremia lactucae* | 20 | Obligate biotrophic | 9767 | 783 | 8.02 |
| *Globisporangium ultimum* | 42 | Necrotrophic | 15322 | 1056 | 6.89 |
| *Hyaloperonospora arabidopsidis* | 18 | Obligate biotrophic | 13982 | 689 | 4.93 |
| *Peronospora effusa* | 19 | Obligate biotrophic | 8603 | 543 | 6.31 |
| *Phytophthora cactorum* | 128 | Hemibiotrophic | 24172 | 2072 | 8.57 |
| *Phytophthora capsici* | 17 | Hemibiotrophic | 19805 | 1293 | 6.53 |
| *Phytophthora cinnamomi* | 6 | Necrotrophic | 26131 | 1862 | 7.13 |
| *Phytophthora fragariae* | 161 | Hemibiotrophic | 41375 | 2809 | 6.79 |
| *Phytophthora infestans* | 92 | Hemibiotrophic | 17797 | 1723 | 9.68 |
| *Phytophthora kernoviae* | 84 | Hemibiotrophic | 10129 | 730 | 7.21 |
| *Phytophthora megakarya* | 57 | Hemibiotrophic | 34804 | 2389 | 6.86 |
| *Phytophthora nicotianae* | 77 | Hemibiotrophic | 17348 | 2059 | 11.87 |
| *Phytophthora parasitica* | 64 | Hemibiotrophic | 27942 | 2223 | 7.96 |
| *Phytophthora rubi* | 93 | Hemibiotrophic | 35524 | 2482 | 6.99 |
| *Phytophthora sojae* | 56 | Hemibiotrophic | 26489 | 2179 | 8.23 |
| *Phytopythium vexans* | 2 | Hemibiotrophic | 11958 | 871 | 7.28 |
| *Plasmopara halstedii* | 19 | Obligate biotrophic | 15469 | 922 | 5.96 |
| *Pythium aphanidermatum* | 1 | Necrotrophic | 12312 | 954 | 7.75 |
| *Pythium insidiosum* | 5 | Other | 14962 | 1088 | 7.27 |
| *Pythium oligandrum* | 45 | Necrotrophic | 14954 | 1556 | 10.41 |
| *Saprolegnia diclina* | 20 | Other | 18229 | 1492 | 8.18 |
| *Saprolegnia parasitica* | 24 | Other | 20121 | 1529 | 7.60 |
| *Thraustotheca clavata* | 11 | Other | 13174 | 789 | 5.99 |
| *Pythium brassicae* | 26 | Necrotrophic | 13005 | 833 | 6.41 |

**Supp. Table 2: Overview of proteins annotated as elicitins.** The table contains all proteins that are annotated with InterPro ID IPR002200. Prediction scores and elicitin classification are included.

**Supp. Table 3: Property summary of elicitin-like proteins from *Albugo*.** Characteristics, expression and prediction results of proteins from *Albugo candida* (Nc2) and *Albugo laibachii* (Nc14) with an annotated elicitin domain. All predictions were done on the accession sequences. Expression refers to infection stages (Early: five days after infection; middle: eight days, late: 14 days). Apoplast proteomics refers to whether or not the proteins have been found in previously done proteomics of apoplastic fluid from *Albugo* infected *Arabidopsis thaliana* at 10 days post infection. The respective dataset for Nc2 and Nc14 has been curated and published[22,64]. aa: amino acid; NT: nucleotide; pos.: position; kDa: kilodalton.

| Name | Accession NCBI | Accession UniProt | Strain | Secretion signal [No. of aa] | Elicitin domain [start – end pos.] | Length [No. of aa] | Mol. Weight [kDa] | Expression | Apoplast proteomics | Mutation compared to accession | Total disorder score | Amyloid prediction |
|---|---|---|---|---|---|---|---|---|---|---|---|---|
| E201 | CCI46865.1 | A0A024GKK9 | Nc2 | 17 | 20-104 | 198 | 19.31 | Late | - | No | 0.298 | 0.965 |
| E202 | CCI47317.1 | A0A024GLV4 | Nc2 | 19 | 82-172 | 229 | 22.78 | Early, late | - | Yes (15 nt changes, 11 aa changes) | 0.490 | 0.958 |
| E203 | CCI47336.1 | A0A024GKD2 | Nc2 | 0 | 4-84 | 141 | 14.82 | Late | - | No | 0.220 | 0.831 |
| E204 | CCI47335.1 | A0A024GLE8 | Nc2 | 0 | 57-147 | 204 | 21.4 | - | - | Yes varying | 0.363 | 0.839 |
| E205 | CCI43375.1 | A0A024GA39 | Nc2 | 19 | 36-126 | 176 | 16.41 | - | - | No | 0.127 | 0.971 |
| E206 | CCI47456.1 | A0A024GKR4 | Nc2 | 19 | 68-158 | 233 | 22.5 | Early, late | Yes | No | 0.514 | 0.965 |
| E207 | CCI42312.1 | A0A024G6C9 | Nc2 | 17 | 18-107 | 165 | 16.34 | Late | - | No | 0.000 | 0.27 |
| E208 | CCI47337.1 | A0A024GLX6 | Nc2 | 0 | 33-123 | 180 | 18.73 | Early, late | - | Yes (5 nt changes, 3 aa changes) | 0.289 | 0.791 |
| E209 | CCI43377.1 | A0A024GAP6 | Nc2 | 20 | 31-120 | 161 | 15.25 | Early, late | - | No | 0.128 | 0.968 |
| E210 | CCI48879.1 | A0A024GQ10 | Nc2 | 19 | 78-168 | 221 | 21.99 | - | - | - | 0.450 | 0.956 |
| E211 | CCI42313.1 | A0A024G6L5 | Nc2 | 30 | 31-120 | 178 | 16.34 | Early, late | - | No | 0.000 | 0.27 |
| E212 | CCI46414.1 | A0A024GJD3 | Nc2 | 22 | 41-107 | 201 | 19.48 | Late | Yes | No | 0.352 | 0.829 |
| E213 | CCI47333.1 | A0A024GKN8 | Nc2 | 19 | 82-172 | 229 | 22.03 | - | - | - | 0.348 | 0.956 |
| E214 | CCI40614.1 | A0A024G1G5 | Nc2 | 0 | 25-118 | 191 | 21.41 | - | - | No | 0.157 | 0.286 |
| E215 | CCI48873.1 | A0A024GR98 | Nc2 | 19 | 82-172 | 229 | 22.55 | Early, late | - | No | 0.367 | 0.978 |
| E216 | CCI47323.1 | A0A024GKM7 | Nc2 | 19 | 86-176 | 233 | 23.33 | Early, late | - | Yes varying | 0.402 | 0.975 |
| E217 | CCI50205.1 | A0A024GTR7 | Nc2 | 0 | 36-109 | 141 | 15.84 | Early, late | - | No | 0.000 | 0.381 |
| E1401 | CCA24552.1 | F0WT64 | Nc14 | 17 | 25-120 | 198 | 19.24 | Middle, late | - | No | 0.387 | 0.959 |
| E1402 | CCA28288.1 | F0X2V9 | Nc14 | 0 | 47-137 | 193 | 20.3 | Middle, late | - | No | 0.404 | 0.798 |
| E1403 | CCA20397.1 | F0WGN3 | Nc14 | 0 | 60-149 | 186 | 20.12 | Middle, late | - | No | 0.000 | 0.092 |
| E1404 | CCA20395.1 | F0WGN1 | Nc14 | 21 | 34-122 | 176 | 16.42 | Middle, late | - | No | 0.187 | 0.979 |
| E1405 | CCA19117.1 | F0WD45 | Nc14 | 15 | 30-105 | 283 | 27.96 | - | Yes | - | 0.608 | 0.938 |
| E1406 | CCA16859.1 | F0W6W4 | Nc14 | 47 | 75-148 | 172 | 14.02 | Middle, late | - | No | 0.000 | 0.79 |
| E1407 | CCA14289.1 | F0VZR8 | Nc14 | 23 | 25-120 | 191 | 18.55 | Middle, late | - | No | 0.167 | 0.914 |
| E1408 | CCA17413.1 | F0W8F8 | Nc14 | 0 | 24-115 | 201 | 22.22 | Middle, late | Yes | Yes partial | 0.318 | 0.064 |
| E1409 | CCA16823.1 | F0W6S8 | Nc14 | 17 | 18-107 | 166 | 16.6 | Middle, late | - | No | 0.000 | 0.369 |

**Supp. Table 4: Primer sequences for *Albugo* elicitin-like proteins.** Primers for amplification of elicitin-like candidates from *Albugo candida* and *Albugo laibachii* for subsequent cloning into the curli-dependent amyloid generator (C-DAG) system[65]. If a secretion signal was predicted for a candidate, it was removed from the sequence before designing the primers. The primers were also used for analyzing expressions at different timepoints.

| Elicitin name | Accession (NCBI) | Forward primer C-DAG cloning from 5' to 3' end | Reverse primer C-DAG cloning from 5' to 3' end | Secretion-signal |
|---|---|---|---|---|
| E201 | CCI46865.1 | TATAGCGGCCGCAGGTGTCTGCAAC | TATATCTAGATTACACAGATACAAA | yes |
| E202 | CCI47317.1 | TATAGCGGCCGCACAAGACGCTCCA | TATATCTAGATTAGTACGAAGCTAA | yes |

| | | | | |
|---|---|---|---|---|
| E203 | CCI47336.1 | TATAGCGGCCGCAGGAATCAATCAG | TATATCTAGATTACAGCAGAGCTAA | no |
| E204 | CCI47335.1 | TATAGCGGCCGCAAGCACAACAAAT | TATATCTAGATTACTGCAGGGCTAG | no |
| E205 | CCI43375.1 | TATAGCGGCCGCACAAACTGCTACA | TATATCTAGATTAATTCAAAGCTAA | yes |
| E206 | CCI47456.1 | TATAGCGGCCGCACAAGGATCTCCA | TATATCTAGATTACTGAATGGCTAT | yes |
| E207 | CCI42312.1 | TATAGCGGCCGCAAAACATTGCCCG | TATATCTAGATTAATTGGGTAGATA | yes |
| E208 | CCI47337.1 | TATAGCGGCCGCACCAAATACCATG | TATATCTAGATTACTGCAGGGCCAA | no |
| E209 | CCI43377.1 | TATAGCGGCCGCAAATGACACCACC | TATATCTAGATTACCAAAGTAACGC | yes |
| E210 | CCI48879.1 | TATAGCGGCCGCACAAGACGCTCCA | TATATCTAGATTAGTACGAAGCCTT | yes |
| E211 | CCI42313.1 | TATAGCGGCCGCAAAACATTGCCCG | TATATCTAGATTAATTGGGTAGATA | yes |
| E212 | CCI46414.1 | TATAGCGGCCGCAAATTATGCGTGC | TATATCTAGATTACACCGTTGCGAT | yes |
| E213 | CCI47333.1 | TATAGCGGCCGCACAAAACATTCCA | TATATCTAGATTACTGCAGGGCGTAA | yes |
| E214 | CCI40614.1 | TATAGCGGCCGCAGCAAAGTACTGT | TATATCTAGATTAATCTTGTCCTTC | no |
| E215 | CCI48873.1 | TATAGCGGCCGCACAAGATGCTCCA | TATATCTAGATTAGTACGAGGCCAG | yes |
| E216 | CCI47323.1 | TATAGCGGCCGCACAAGACGCTCCA | TATATCTAGATTAGTACGAAGCCAA | yes |
| E217 | CCI50205.1 | TATAGCGGCCGCAGTTTCCCATTTC | TATATCTAGATTATGTATGTGTCAA | no |
| E1401 | CCA24552.1 | TATAGCGGCCGCAGATGTCTGCAAC | TATATCTAGATTACACGAATTCGAG | yes |
| E1402 | CCA28288.1 | TATAGCGGCCGCAGGTGATTCATCC | TATATCTAGATTACTGCATGGCTAA | no |
| E1403 | CCA20397.1 | TATAGCGGCCGCAGAAACTTTCCGC | TATATCTAGATTACAACGACAATGC | no |
| E1404 | CCA20395.1 | TATAGCGGCCGCACAAGGTACTCAA | TATATCTAGATTAATTGAAAGCTAA | yes |
| E1405 | CCA19117.1 | TATAGCGGCCGCAGCCGAAGACAC | TATATCTAGATTACATGAAAAAGAA | yes |
| E1406 | CCA16859.1 | TATAGCGGCCGCAGCTGATCCGCCT | TATATCTAGATTATATACGAGTGCC | yes |
| E1407 | CCA14289.1 | TATAGCGGCCGCACAAATCCGATGC | TATATCTAGATTAATCTTGACCTTC | yes |
| E1408 | CCA17413.1 | TATAGCGGCCGCACGAATCACTTCC | TATATCTAGATTACGCTGTTGCGAT | no |
| E1409 | CCA16823.1 | TATAGCGGCCGCAGATCATTGCCCA | TATATCTAGATTAGTTGGAAAGATA | yes |
| E205Core | Partial CCI43375.1 | TATAGCGGCCGCATGCAATGAAAAG | TATATCTAGATTAGCACTGCTGTAC | - |
| E215Core | Partial CCI48873.1 | TATAGCGGCCGCATGCAATGACACT | TATATCTAGATTAGCACAAACCGGT | - |
| E206Core | Partial CCI47456.1 | TATAGCGGCCGCATGCAGTGATAAG | TATATCTAGATTAGCAGCCATTGAC | - |
| E212Core | Partial CCI46414.1 | TATAGCGGCCGCAAATTATGCGTGC | TATATCTAGATTAGATTAGCCATCC | yes |
| E212IDR | Partial CCI46414.1 | TATAGCGGCCGCATCAACGTGGTTG | TATATCTAGATTACACCGTTGCGAT | - |

**Supp. Table 5: Primer sequences for INF1 constructs.** Primers for amplification of elicitin candidate INF1 from *Phytophthora infestans* for subsequent cloning into the PURExpress® system. The secretion signal was removed from the sequence before designing the primers.

| Elicitin name | Accession (UniProt) | Forward primer 5' to 3' end | Reverse primer 5' to 3' end | Secretion-signal predicted |
|---|---|---|---|---|
| INF1 | Q01905 | GGAGATATACATATGACCACGTGCACCACCTCG | AACCTTACTCGACTATCATAGCGACGCACACGTAGACG | yes |
| INF1-E212IDR | Q01905 | AAGGAGATATACATATGACCACGTGCACCACCTC | AACCACGTTGACATAAGCGACGCACACGTAGACG | yes |

**Supp. Table 6: Primer sequences for PURExpress system.** Primers for amplification of elicitin-like candidates from *Albugo candida* and *Albugo laibachii* for subsequent cloning into the PURExpress® system. If a secretion signal was predicted for a candidate, it was removed from the sequence before designing the primers. NCBI: National Center for Biological Information.

| Elicitin name | Accession (NCBI) | Forward primer PURExpress cloning from 5' to 3' end | Reverse primer PURExpress cloning from 5' to 3' end | Secretion-signal predicted |
|---|---|---|---|---|
| E212 | CCI46414.1 | AAGGAGATATACATATGAATTATGCGTGCGATATTCCTCAAATCC | GTTAACCTTACTCGACTACACCGTTGCGATGGAAACA | yes |
| E212Core | Partial CCI46414.1 | AAGGAGATATACATATGAATTATGCGTGCGATATT | GTTAACCTTACTCGACTAGATTAGCCATCCATACGA | yes |
| E212IDR | Partial CCI46414.1 | AAGGAGATATACATATGTCAACGTGGTTGATGGGA | GTTAACCTTACTCGACTACACCGTTGCGATGGAAAC | - |
| E206 | CCI47456.1 | AAGGAGATATACATATGCAAGGATCTCCATCCGTCACC | GTTAACCTTACTCGACTACTGAATGGCTATCACGAAGACAGC | yes |
| E206Core | Partial CCI47456.1 | AAGGAGATATACATATGTGCAGTGATAAGGTCGCGC | GTTAACCTTACTCGACTAGCAGCCATTGACAAACATCTTCG | - |
| E207 | CCI42312.1 | GGAGATATACATATGAAACATTGCCCGAGCTCTG | AACCTTACTCGACTAATTGGGTAGATACAATATCTCCGAG | yes |

**Supp. Table 7: LC–MS/MS analysis.** Monitored transitions after targeted LC–MS/MS analysis on the aggregation band of E212 expressed via PURExpress system and extracted after SDS PAGE.

| Q1 (m/z) | Q3 (m/z) | peptide | CE (Volts) |
|---|---|---|---|
| 201.799 | 216.1343 | Nc2_E212.SKIQK.+3b2.light | 17 |
| 201.799 | 229.1421 | Nc2_E212.SKIQK.+3b4+2.light | 17 |
| 201.799 | 258.6788 | Nc2_E212.SKIQK.+3y4+2.light | 15 |
| 201.799 | 275.1714 | Nc2_E212.SKIQK.+3y2.light | 15 |
| 201.799 | 329.2183 | Nc2_E212.SKIQK.+3b3.light | 15 |
| 201.799 | 388.2554 | Nc2_E212.SKIQK.+3y3.light | 15 |
| 201.799 | 457.2769 | Nc2_E212.SKIQK.+3b4.light | 15 |
| 201.799 | 516.3504 | Nc2_E212.SKIQK.+3y4.light | 17 |
| 221.4379 | 230.0897 | Nc2_E212.DEDVGK.+3b4+2.light | 20 |
| 221.4379 | 245.0768 | Nc2_E212.DEDVGK.+3b2.light | 20 |
| 221.4379 | 258.6005 | Nc2_E212.DEDVGK.+3b5+2.light | 20 |
| 221.4379 | 274.1397 | Nc2_E212.DEDVGK.+3y5+2.light | 20 |
| 221.4379 | 303.2027 | Nc2_E212.DEDVGK.+3y3.light | 20 |
| 221.4379 | 360.1038 | Nc2_E212.DEDVGK.+3b3.light | 20 |
| 221.4379 | 418.2296 | Nc2_E212.DEDVGK.+3y4.light | 20 |
| 221.4379 | 459.1722 | Nc2_E212.DEDVGK.+3b4.light | 20 |
| 221.4379 | 547.2722 | Nc2_E212.DEDVGK.+3y5.light | 20 |
| 293.1469 | 317.1819 | Nc2_E212.DEDVGKSK.+3y6+2.light | 22 |
| 293.1469 | 322.6479 | Nc2_E212.DEDVGKSK.+3b6+2.light | 22 |
| 293.1469 | 360.1038 | Nc2_E212.DEDVGKSK.+3b3.light | 22 |
| 293.1469 | 362.2398 | Nc2_E212.DEDVGKSK.+3y3.light | 22 |
| 293.1469 | 366.164 | Nc2_E212.DEDVGKSK.+3b7+2.light | 22 |
| 293.1469 | 381.7032 | Nc2_E212.DEDVGKSK.+3y7+2.light | 22 |
| 293.1469 | 419.2613 | Nc2_E212.DEDVGKSK.+3y4.light | 22 |
| 293.1469 | 459.1722 | Nc2_E212.DEDVGKSK.+3b4.light | 22 |
| 293.1469 | 516.1936 | Nc2_E212.DEDVGKSK.+3b5.light | 22 |
| 293.1469 | 518.3297 | Nc2_E212.DEDVGKSK.+3y5.light | 22 |
| 302.1949 | 329.2183 | Nc2_E212.SKIQK.+2b3.light | 17 |

| | | | |
|---|---|---|---|
| 302.1949 | 388.2554 | Nc2_E212.SKIQK.+2y3.light | 17 |
| 302.1949 | 457.2769 | Nc2_E212.SKIQK.+2b4.light | 17 |
| 302.1949 | 516.3504 | Nc2_E212.SKIQK.+2y4.light | 17 |
| 331.6532 | 360.1038 | Nc2_E212.DEDVGK.+2b3.light | 20 |
| 331.6532 | 418.2296 | Nc2_E212.DEDVGK.+2y4.light | 20 |
| 331.6532 | 459.1722 | Nc2_E212.DEDVGK.+2b4.light | 20 |
| 331.6532 | 516.1936 | Nc2_E212.DEDVGK.+2b5.light | 20 |
| 331.6532 | 547.2722 | Nc2_E212.DEDVGK.+2y5.light | 20 |
| 382.1889 | 385.6714 | Nc2_E212.SGYDIFSLDK.+3b7+2.light | 25 |
| 382.1889 | 419.2213 | Nc2_E212.SGYDIFSLDK.+3y7+2.light | 25 |
| 382.1889 | 423.151 | Nc2_E212.SGYDIFSLDK.+3b4.light | 25 |
| 382.1889 | 442.2134 | Nc2_E212.SGYDIFSLDK.+3b8+2.light | 25 |
| 382.1889 | 462.2558 | Nc2_E212.SGYDIFSLDK.+3y4.light | 25 |
| 382.1889 | 499.7269 | Nc2_E212.SGYDIFSLDK.+3b9+2.light | 25 |
| 382.1889 | 500.7529 | Nc2_E212.SGYDIFSLDK.+3y8+2.light | 25 |
| 382.1889 | 529.2637 | Nc2_E212.SGYDIFSLDK.+3y9+2.light | 25 |
| 382.1889 | 536.2351 | Nc2_E212.SGYDIFSLDK.+3b5.light | 25 |
| 382.1889 | 609.3243 | Nc2_E212.SGYDIFSLDK.+3y5.light | 25 |
| 382.1889 | 683.3035 | Nc2_E212.SGYDIFSLDK.+3b6.light | 25 |
| 382.1889 | 722.4083 | Nc2_E212.SGYDIFSLDK.+3y6.light | 25 |
| 439.2167 | 459.1722 | Nc2_E212.DEDVGKSK.+2b4.light | 22 |
| 439.2167 | 516.1936 | Nc2_E212.DEDVGKSK.+2b5.light | 22 |
| 439.2167 | 518.3297 | Nc2_E212.DEDVGKSK.+2y5.light | 22 |
| 439.2167 | 633.3566 | Nc2_E212.DEDVGKSK.+2y6.light | 22 |
| 439.2167 | 644.2886 | Nc2_E212.DEDVGKSK.+2b6.light | 22 |
| 439.2167 | 762.3992 | Nc2_E212.DEDVGKSK.+2y7.light | 22 |
| 483.5648 | 489.278 | Nc2_E212.EC[CAM]TDVLNQINSR.+3y4.light | 25 |
| 483.5648 | 506.1551 | Nc2_E212.EC[CAM]TDVLNQINSR.+3b4.light | 25 |
| 483.5648 | 529.7831 | Nc2_E212.EC[CAM]TDVLNQINSR.+3y9+2.light | 25 |
| 483.5648 | 537.2502 | Nc2_E212.EC[CAM]TDVLNQINSR.+3b9+2.light | 25 |
| 483.5648 | 580.3069 | Nc2_E212.EC[CAM]TDVLNQINSR.+3y10+2.light | 25 |
| 483.5648 | 594.2717 | Nc2_E212.EC[CAM]TDVLNQINSR.+3b10+2.light | 25 |
| 483.5648 | 605.2236 | Nc2_E212.EC[CAM]TDVLNQINSR.+3b5.light | 25 |
| 483.5648 | 617.3365 | Nc2_E212.EC[CAM]TDVLNQINSR.+3y5.light | 25 |
| 483.5648 | 637.7877 | Nc2_E212.EC[CAM]TDVLNQINSR.+3b11+2.light | 25 |
| 483.5648 | 660.3223 | Nc2_E212.EC[CAM]TDVLNQINSR.+3y11+2.light | 25 |
| 483.5648 | 718.3076 | Nc2_E212.EC[CAM]TDVLNQINSR.+3b6.light | 25 |
| 483.5648 | 731.3795 | Nc2_E212.EC[CAM]TDVLNQINSR.+3y6.light | 25 |
| 500.2463 | 504.2489 | Nc2_E212.EQKPAHASASTESR.+3b10+2.light | 28 |
| 500.2463 | 508.7414 | Nc2_E212.EQKPAHASASTESR.+3y10+2.light | 28 |
| 500.2463 | 554.2933 | Nc2_E212.EQKPAHASASTESR.+3b5.light | 28 |
| 500.2463 | 554.7727 | Nc2_E212.EQKPAHASASTESR.+3b11+2.light | 28 |
| 500.2463 | 557.2678 | Nc2_E212.EQKPAHASASTESR.+3y11+2.light | 28 |
| 500.2463 | 579.2733 | Nc2_E212.EQKPAHASASTESR.+3y5.light | 28 |
| 500.2463 | 619.294 | Nc2_E212.EQKPAHASASTESR.+3b12+2.light | 28 |
| 500.2463 | 621.3153 | Nc2_E212.EQKPAHASASTESR.+3y12+2.light | 28 |
| 500.2463 | 650.3104 | Nc2_E212.EQKPAHASASTESR.+3y6.light | 28 |
| 500.2463 | 691.3522 | Nc2_E212.EQKPAHASASTESR.+3b6.light | 28 |
| 500.2463 | 737.3424 | Nc2_E212.EQKPAHASASTESR.+3y7.light | 28 |
| 500.2463 | 762.3893 | Nc2_E212.EQKPAHASASTESR.+3b7.light | 28 |
| 542.9447 | 557.2678 | Nc2_E212.KEQKPAHASASTESR.+3y11+2.light | 28 |
| 542.9447 | 568.2964 | Nc2_E212.KEQKPAHASASTESR.+3b11+2.light | 28 |
| 542.9447 | 579.2733 | Nc2_E212.KEQKPAHASASTESR.+3y5.light | 28 |
| 542.9447 | 611.3511 | Nc2_E212.KEQKPAHASASTESR.+3b5.light | 28 |
| 542.9447 | 618.8202 | Nc2_E212.KEQKPAHASASTESR.+3b12+2.light | 28 |
| 542.9447 | 621.3153 | Nc2_E212.KEQKPAHASASTESR.+3y12+2.light | 28 |
| 542.9447 | 650.3104 | Nc2_E212.KEQKPAHASASTESR.+3y6.light | 28 |
| 542.9447 | 682.3883 | Nc2_E212.KEQKPAHASASTESR.+3b6.light | 28 |
| 542.9447 | 683.3415 | Nc2_E212.KEQKPAHASASTESR.+3b13+2.light | 28 |
| 542.9447 | 685.3446 | Nc2_E212.KEQKPAHASASTESR.+3y13+2.light | 28 |

| | | | |
|---|---|---|---|
| 542.9447 | 737.3424 | Nc2_E212.KEQKPAHASASTESR.+3y7.light | 28 |
| 542.9447 | 819.4472 | Nc2_E212.KEQKPAHASASTESR.+3b7.light | 28 |
| 567.2848 | 598.8227 | Nc2_E212.YPNEEQVQVLSHTR.+3y10+2.light | 25.2 |
| 567.2848 | 600.7984 | Nc2_E212.YPNEEQVQVLSHTR.+3b10+2.light | 25.2 |
| 567.2848 | 613.3416 | Nc2_E212.YPNEEQVQVLSHTR.+3y5.light | 25.2 |
| 567.2848 | 633.2515 | Nc2_E212.YPNEEQVQVLSHTR.+3b5.light | 25.2 |
| 567.2848 | 644.3144 | Nc2_E212.YPNEEQVQVLSHTR.+3b11+2.light | 25.2 |
| 567.2848 | 663.344 | Nc2_E212.YPNEEQVQVLSHTR.+3y11+2.light | 25.2 |
| 567.2848 | 712.41 | Nc2_E212.YPNEEQVQVLSHTR.+3y6.light | 25.2 |
| 567.2848 | 712.8439 | Nc2_E212.YPNEEQVQVLSHTR.+3b12+2.light | 25.2 |
| 567.2848 | 720.3655 | Nc2_E212.YPNEEQVQVLSHTR.+3y12+2.light | 25.2 |
| 567.2848 | 761.3101 | Nc2_E212.YPNEEQVQVLSHTR.+3b6.light | 25.2 |
| 567.2848 | 840.4686 | Nc2_E212.YPNEEQVQVLSHTR.+3y7.light | 25.2 |
| 567.2848 | 860.3785 | Nc2_E212.YPNEEQVQVLSHTR.+3b7.light | 25.2 |
| 572.7797 | 609.3243 | Nc2_E212.SGYDIFSLDK.+2y5.light | 27.1 |
| 572.7797 | 683.3035 | Nc2_E212.SGYDIFSLDK.+2b6.light | 27.1 |
| 572.7797 | 722.4083 | Nc2_E212.SGYDIFSLDK.+2y6.light | 27.1 |
| 572.7797 | 770.3355 | Nc2_E212.SGYDIFSLDK.+2b7.light | 27.1 |
| 572.7797 | 837.4353 | Nc2_E212.SGYDIFSLDK.+2y7.light | 27.1 |
| 572.7797 | 883.4196 | Nc2_E212.SGYDIFSLDK.+2b8.light | 27.1 |
| 724.8435 | 731.3795 | Nc2_E212.EC[CAM]TDVLNQINSR.+2y6.light | 34.5 |
| 724.8435 | 832.3505 | Nc2_E212.EC[CAM]TDVLNQINSR.+2b7.light | 34.5 |
| 724.8435 | 844.4635 | Nc2_E212.EC[CAM]TDVLNQINSR.+2y7.light | 34.5 |
| 724.8435 | 943.532 | Nc2_E212.EC[CAM]TDVLNQINSR.+2y8.light | 34.5 |
| 724.8435 | 960.4091 | Nc2_E212.EC[CAM]TDVLNQINSR.+2b8.light | 34.5 |
| 749.8659 | 762.3893 | Nc2_E212.EQKPAHASASTESR.+2b7.light | 35.7 |
| 749.8659 | 808.3795 | Nc2_E212.EQKPAHASASTESR.+2y8.light | 35.7 |
| 749.8659 | 849.4213 | Nc2_E212.EQKPAHASASTESR.+2b8.light | 35.7 |
| 749.8659 | 920.4585 | Nc2_E212.EQKPAHASASTESR.+2b9.light | 35.7 |
| 749.8659 | 945.4384 | Nc2_E212.EQKPAHASASTESR.+2y9.light | 35.7 |
| 804.7485 | 830.4609 | Nc2_E212.ATSVC[CAM]FSFVVGGFFAFVVSIATV.+3y16+2.light | 36.6 |
| 804.7485 | 835.4924 | Nc2_E212.ATSVC[CAM]FSFVVGGFFAFVVSIATV.+3y8.light | 36.6 |
| 804.7485 | 862.9107 | Nc2_E212.ATSVC[CAM]FSFVVGGFFAFVVSIATV.+3b16+2.light | 36.6 |
| 804.7485 | 873.9769 | Nc2_E212.ATSVC[CAM]FSFVVGGFFAFVVSIATV.+3y17+2.light | 36.6 |
| 804.7485 | 900.392 | Nc2_E212.ATSVC[CAM]FSFVVGGFFAFVVSIATV.+3b8.light | 36.6 |
| 804.7485 | 906.5295 | Nc2_E212.ATSVC[CAM]FSFVVGGFFAFVVSIATV.+3y9.light | 36.6 |
| 804.7485 | 912.4449 | Nc2_E212.ATSVC[CAM]FSFVVGGFFAFVVSIATV.+3b17+2.light | 36.6 |
| 804.7485 | 947.5111 | Nc2_E212.ATSVC[CAM]FSFVVGGFFAFVVSIATV.+3y18+2.light | 36.6 |
| 804.7485 | 961.9791 | Nc2_E212.ATSVC[CAM]FSFVVGGFFAFVVSIATV.+3b18+2.light | 36.6 |
| 804.7485 | 999.4604 | Nc2_E212.ATSVC[CAM]FSFVVGGFFAFVVSIATV.+3b9.light | 36.6 |
| 850.4236 | 860.3785 | Nc2_E212.YPNEEQVQVLSHTR.+2b7.light | 40.7 |
| 850.4236 | 939.537 | Nc2_E212.YPNEEQVQVLSHTR.+2y8.light | 40.7 |
| 850.4236 | 988.4371 | Nc2_E212.YPNEEQVQVLSHTR.+2b8.light | 40.7 |
| 421.75 | 472.3 | Trypsin 421 472 | 20 |
| 421.75 | 571.4 | Trypsin 421 571 | 20 |

**Supp. Table 8: Overview of plant infiltration experiments in *Arabidopsis thaliana*.** Sets and rounds were conducted independently. Plants were grown specifically for the individual rounds, and protein was produced fresh for each infiltration.

| Set | Round | Plant | Count of plants per treatment | Count of infiltrated leaves | Day 1 Infiltration | Day 2 Infiltration |
|---|---|---|---|---|---|---|
| 1 | 1 | *Arabidopsis thaliana* WS-0 | 4 | 20 | Buffer | Buffer |
| 1 | 1 | *Arabidopsis thaliana* WS-0 | 4 | 20 | E212 | E212 |
| 1 | 1 | *Arabidopsis thaliana* WS-0 | 4 | 20 | E212Core | E212Core |
| 1 | 1 | *Arabidopsis thaliana* WS-0 | 4 | 20 | E212IDR | E212IDR |
| 1 | 2 | *Arabidopsis thaliana* WS-0 | 4 | 20 | Buffer | Buffer |

| | | | | | | |
|---|---|---|---|---|---|---|
| 1 | 2 | *Arabidopsis thaliana* WS-0 | 4 | 20 | E212 | E212 |
| 1 | 2 | *Arabidopsis thaliana* WS-0 | 4 | 20 | E212Core | E212Core |
| 1 | 2 | *Arabidopsis thaliana* WS-0 | 4 | 20 | E212IDR | E212IDR |
| 1 | 3 | *Arabidopsis thaliana* WS-0 | 4 | 20 | Buffer | Buffer |
| 1 | 3 | *Arabidopsis thaliana* WS-0 | 4 | 20 | E212 | E212 |
| 1 | 3 | *Arabidopsis thaliana* WS-0 | 4 | 20 | E212Core | E212Core |
| 1 | 3 | *Arabidopsis thaliana* WS-0 | 4 | 20 | E212IDR | E212IDR |
| 2 | 1 | *Arabidopsis thaliana* WS-0 | 4 | 20 | Buffer | Buffer |
| 2 | 1 | *Arabidopsis thaliana* WS-0 | 4 | 20 | E206 | E206 |
| 2 | 1 | *Arabidopsis thaliana* WS-0 | 4 | 20 | E206Core | E206Core |
| 2 | 1 | *Arabidopsis thaliana* WS-0 | 4 | 20 | E207 | E207 |
| 2 | 2 | *Arabidopsis thaliana* WS-0 | 4 | 20 | Buffer | Buffer |
| 2 | 2 | *Arabidopsis thaliana* WS-0 | 4 | 20 | E206 | E206 |
| 2 | 2 | *Arabidopsis thaliana* WS-0 | 4 | 20 | E206Core | E206Core |
| 2 | 2 | *Arabidopsis thaliana* WS-0 | 4 | 20 | E207 | E207 |
| 2 | 3 | *Arabidopsis thaliana* WS-0 | 4 | 20 | Buffer | Buffer |
| 2 | 3 | *Arabidopsis thaliana* WS-0 | 4 | 20 | E206 | E206 |
| 2 | 3 | *Arabidopsis thaliana* WS-0 | 4 | 20 | E206Core | E206Core |
| 2 | 3 | *Arabidopsis thaliana* WS-0 | 4 | 20 | E207 | E207 |
| 3 | 1 | *Arabidopsis thaliana* Col-0 | 4 | 20 | E212Core | E212Core |
| 3 | 1 | *Arabidopsis thaliana* Col-0 bak1-3 | 4 | 20 | E212Core | E212Core |
| 3 | 1 | *Arabidopsis thaliana* Col-0 bak1-5 | 4 | 20 | E212Core | E212Core |
| 3 | 2 | *Arabidopsis thaliana* Col-0 | 4 | 20 | E212Core | E212Core |
| 3 | 2 | *Arabidopsis thaliana* Col-0 bak1-3 | 4 | 20 | E212Core | E212Core |
| 3 | 2 | *Arabidopsis thaliana* Col-0 bak1-5 | 4 | 20 | E212Core | E212Core |
| 3 | 3 | *Arabidopsis thaliana* Col-0 | 4 | 20 | E212Core | E212Core |
| 3 | 3 | *Arabidopsis thaliana* Col-0 bak1-3 | 4 | 20 | E212Core | E212Core |
| 3 | 3 | *Arabidopsis thaliana* Col-0 bak1-5 | 4 | 20 | E212Core | E212Core |
| 3 | 3 | *Arabidopsis thaliana* Col-0 | 4 | 20 | Buffer | Buffer |

**Supp. Table 9: Overview of plant infiltration experiments in *Nicotiana benthamiana*.** Sets and rounds were conducted independently. Plants were grown specifically for the individual rounds, and protein was produced fresh for each infiltration. For some treatments in set 3 and 4 different proteins were infiltrated on day 1 and 2.

| Set | Round | Plant | Count of plants per treatment | Day 1 Infiltration | Day 2 Infiltration |
|---|---|---|---|---|---|
| 1 | 1 | *Nicotiana benthamiana* | 3 | INF1 | INF1 |
| 1 | 1 | *Nicotiana benthamiana* | 3 | E206Core | E206Core |
| 1 | 1 | *Nicotiana benthamiana* | 3 | E212Core | E212Core |
| 1 | 1 | *Nicotiana benthamiana* | 3 | Buffer | Buffer |

| | | | | | |
|---|---|---|---|---|---|
| 1 | 2 | *Nicotiana benthamiana* | 3 | INF1 | INF1 |
| 1 | 2 | *Nicotiana benthamiana* | 3 | E206Core | E206Core |
| 1 | 2 | *Nicotiana benthamiana* | 3 | E212Core | E212Core |
| 1 | 2 | *Nicotiana benthamiana* | 3 | Buffer | Buffer |
| 1 | 3 | *Nicotiana benthamiana* | 3 | INF1 | INF1 |
| 1 | 3 | *Nicotiana benthamiana* | 3 | E206Core | E206Core |
| 1 | 3 | *Nicotiana benthamiana* | 3 | E212Core | E212Core |
| 1 | 3 | *Nicotiana benthamiana* | 3 | Buffer | Buffer |
| 2 | 1 | *Nicotiana benthamiana* | 3 | INF1 | INF1 |
| 2 | 1 | *Nicotiana benthamiana* | 3 | INF1-E212IDR | INF1-E212IDR |
| 2 | 1 | *Nicotiana benthamiana* | 3 | Buffer | Buffer |
| 2 | 1 | *Nicotiana benthamiana* | 3 | - | - |
| 2 | 2 | *Nicotiana benthamiana* | 3 | INF1 | INF1 |
| 2 | 2 | *Nicotiana benthamiana* | 3 | INF1-E212IDR | INF1-E212IDR |
| 2 | 2 | *Nicotiana benthamiana* | 3 | Buffer | Buffer |
| 2 | 2 | *Nicotiana benthamiana* | 3 | - | - |
| 2 | 3 | *Nicotiana benthamiana* | 3 | INF1 | INF1 |
| 2 | 3 | *Nicotiana benthamiana* | 3 | INF1-E212IDR | INF1-E212IDR |
| 2 | 3 | *Nicotiana benthamiana* | 3 | Buffer | Buffer |
| 2 | 3 | *Nicotiana benthamiana* | 3 | - | - |
| 3 | 1 | *Nicotiana benthamiana* | 3 | 0.5x INF1 | 0.5x INF1 |
| 3 | 1 | *Nicotiana benthamiana* | 3 | E212IDR | INF1 |
| 3 | 1 | *Nicotiana benthamiana* | 3 | Buffer | INF1 |
| 3 | 1 | *Nicotiana benthamiana* | 3 | Buffer | Buffer |
| 3 | 2 | *Nicotiana benthamiana* | 3 | 0.5x INF1 | 0.5x INF1 |
| 3 | 2 | *Nicotiana benthamiana* | 3 | E212IDR | INF1 |
| 3 | 2 | *Nicotiana benthamiana* | 3 | Buffer | INF1 |
| 3 | 2 | *Nicotiana benthamiana* | 3 | Buffer | Buffer |
| 3 | 3 | *Nicotiana benthamiana* | 3 | 0.5x INF1 | 0.5x INF1 |
| 3 | 3 | *Nicotiana benthamiana* | 3 | E212IDR | INF1 |
| 3 | 3 | *Nicotiana benthamiana* | 3 | Buffer | INF1 |
| 3 | 3 | *Nicotiana benthamiana* | 3 | Buffer | Buffer |
| 4 | 1 | *Nicotiana benthamiana* | 3 | INF1 | INF1 |
| 4 | 1 | *Nicotiana benthamiana* | 3 | INF1 + E212IDR | INF1 + E212IDR |
| 4 | 1 | *Nicotiana benthamiana* | 3 | INF1-E212IDR | INF1-E212IDR |
| 4 | 1 | *Nicotiana benthamiana* | 3 | E212IDR | E212IDR |
| 4 | 2 | *Nicotiana benthamiana* | 3 | INF1 | INF1 |
| 4 | 2 | *Nicotiana benthamiana* | 3 | INF1 + E212IDR | INF1 + E212IDR |
| 4 | 2 | *Nicotiana benthamiana* | 3 | INF1-E212IDR | INF1-E212IDR |
| 4 | 2 | *Nicotiana benthamiana* | 3 | E212IDR | E212IDR |
| 4 | 3 | *Nicotiana benthamiana* | 3 | INF1 | INF1 |
| 4 | 3 | *Nicotiana benthamiana* | 3 | INF1 + E212IDR | INF1 + E212IDR |
| 4 | 3 | *Nicotiana benthamiana* | 3 | INF1-E212IDR | INF1-E212IDR |
| 4 | 3 | *Nicotiana benthamiana* | 3 | E212IDR | E212IDR |
